# Deciphering the LRRK code: LRRK1 and LRRK2 phosphorylate distinct Rab proteins and are regulated by diverse mechanisms

**DOI:** 10.1101/2020.11.25.397836

**Authors:** Asad U. Malik, Athanasios Karapetsas, Raja S. Nirujogi, Sebastian Mathea, Prosenjit Pal, Pawel Lis, Matthew Taylor, Elena Purlyte, Robert Gourlay, Mark Dorward, Simone Weidlich, Rachel Toth, Nicole K. Polinski, Stefan Knapp, Francesca Tonelli, Dario R Alessi

**Author notes:** Correspondence to Francesca Tonelli or Dario R Alessi.

## Abstract

Much attention has focused on LRRK2, as autosomal dominant missense mutations that enhance its kinase activity cause inherited Parkinson’s disease. LRRK2 regulates biology by phosphorylating a subset of Rab GTPases including Rab8A and Rab10 within its effector binding motif. In this study we explore whether LRRK1, a less studied homologue of LRRK2 that regulates growth factor receptor trafficking and osteoclast biology might also phosphorylate Rab proteins. Using mass spectrometry, we found that the endogenous Rab7A protein, phosphorylated at Ser72 was most impacted by LRRK1 knock-out. This residue is not phosphorylated by LRRK2 but lies at the equivalent site targeted by LRRK2 on Rab8A and Rab10. Accordingly, recombinant LRRK1 efficiently phosphorylated Rab7A at Ser72, but not Rab8A or Rab10. Employing a novel phospho-specific antibody, we found that phorbol ester stimulation of mouse embryonic fibroblasts markedly enhanced phosphorylation of Rab7A at Ser72 via LRRK1. We identify two LRRK1 mutations (K746G and I1412T), equivalent to the LRRK2 R1441G and I2020T Parkinson’s mutations, that enhance LRRK1 mediated phosphorylation of Rab7A. We demonstrate that two regulators of LRRK2 namely Rab29 and VPS35[D620N], do not influence LRRK1. Widely used LRRK2 inhibitors do not inhibit LRRK1, but we identify a promiscuous Type-2 tyrosine kinase inhibitor termed GZD-824 that inhibits both LRRK1 and LRRK2. Finally, we show that interaction of Rab7A with its effector RILP is not affected by high stoichiometry LRRK1 phosphorylation. Altogether, these finding reinforce the idea that the LRRK enzymes have evolved as major regulators of Rab biology.

## Introduction

Mammals possess two homologues of the leucine-rich repeat protein kinase termed LRRK1 and LRRK2 [1]. These are large multidomain enzymes possessing an N-terminal ankyrin motif, followed by leucine-rich repeats, a Roco module comprising a Roc GTPase and a COR dimerization motif, a protein kinase and a C-terminal WD40 domain [2]. LRRK1 is slightly smaller than LRRK2 due to the lack of an N-terminal armadillo domain. Comparative low resolution cryo-EM analysis suggested that LRRK1 and LRRK2 form homodimers possessing similar overall shapes [3]. Both enzymes bind and hydrolyze GTP through their Roc GTPase domains [4, 5].

Despite overall domain similarity, LRRK1 and LRRK2 appear to possess distinct physiological functions. Genetic evidence in humans, points towards LRRK1 regulating bone biology and LRRK2 contributing to Parkinson’s disease. Autosomal recessive variants that induce frameshift or truncating mutations within the C-terminal domain of LRRK1, that are likely loss of function, cause a severe metabolic bone disorder termed osteosclerotic metaphyseal dysplasia (OSMD) [6-9]. This is a unique form of osteopetrosis characterized by severe osteosclerosis localized to the metaphyses of long bones [10]. LRRK1 knock-out mice also display severe osteopetrosis that is not observed in LRRK2 deficient animals [11, 12]. In contrast, autosomal dominant missense mutations located within the GTPase or kinase catalytic domains that hyperactivate LRRK2 protein kinase activity, are one of the most common causes of inherited Parkinson’s disease [13, 14].

LRRK2 phosphorylates a subgroup of Rab proteins (Rab3A/B/C/D, Rab8A/B, Rab10, Rab12, Rab29, Rab35, and Rab43) at a conserved Thr/Ser residue (Thr73 for Rab10), located within the effector binding switch-II motif [15-17]. As mentioned above, Parkinson’s pathogenic mutations increase LRRK2 protein kinase activity thereby stimulating Rab protein phosphorylation [15, 16, 18]. In turn, this promotes their interaction with a group of effectors including RILPL1, RILPL2, JIP3 and JIP4 that bind preferentially to LRRK2 phosphorylated Rab8 and Rab10 through a phosphorylation-specific effector binding RH2 domain [15, 19]. LRRK2 phosphorylated Rab8A and Rab10, in complex with RILPL1, inhibit the formation of primary cilia that are implicated in controlling a Sonic hedgehog-driven neuroprotective pathway that could provide a mechanism by which LRRK2 is linked to Parkinson’s disease [20]. Other research has revealed that other components genetically linked to Parkinson’s disease including Rab29 [21, 22] and VPS35 [23] also regulate phosphorylation of Rab proteins via LRRK2.

Co-immunoprecipitation studies revealed that LRRK1 interacts with numerous components of the EGF Receptor signalling pathway [24-26]. EGF stimulation promotes accumulation of LRRK1 at the endosome and this has been proposed to control trafficking of EGF receptors from the cell surface to intracellular endocytic compartments [24, 26]. It has also been reported that the EGF receptor tyrosine kinase inhibits LRRK1 protein kinase activity by directly phosphorylating Tyr971, a residue that is located within the Roco dimerization COR domain [27]. Overexpression of LRRK1 in cells was also shown to enhance dynein-mediated transport of EGF receptor-containing endosomes toward the perinuclear region by promoting phosphorylation of GTP-bound Rab7A on Ser72 and enhancing its interaction with an effector termed RILP [28].

Other work has suggested that LRRK1 is recruited to the autolysosome during autophagy, where it activates the Rab7 GTPase-activating protein termed TBC1D2, thereby switching off Rab7A function [29]. LRRK1 has also been associated with controlling microtubule dynein-driven transport (perhaps via RILP) [30], mitotic spindle orientation, [31] as well as in NF-κB signaling in B cells and the humoral immune response [32]. Various LRRK1 substrates have been proposed including CLIP-170 (Thr1384) [30], CDK5RAP2 (Thr 102 and Ser 140) [31], RAC1/Cdc42 small GTPases (Ser71) [33], L-plastin (Ser 5) [34], in addition to Rab7A [28]. To our knowledge the phosphorylation of these endogenous proteins by LRRK1 has not yet been independently validated. Given that Rab proteins are the key substrates of LRRK2, we sought to explore whether LRRK1 could regulate the phosphorylation of Rab proteins. Our data suggest that a key substrate for LRRK1 is Rab7A that is phosphorylated at Ser72. Moreover, LRRK1 does not appear to phosphorylate Rab8A or Rab10, that are key substrates for LRRK2. Our data also reveal that LRRK1 and LRRK2 are regulated by distinct upstream mechanisms.

## Materials and Methods

### Reagents

[γ-^32^P]ATP was purchased from PerkinElmer. HG-10-102-01, MLi-2, GSK2578215A, LRRK2-IN1, MRT67307, GDC0941, AG1478, BID1870, H-89, PD0325901, UO1216, Rapamycin and Phos-tag acrylamide were all synthesized by Natalia Shpiro (University of Dundee). Phos-tag acrylamide stocks are stored in aliquots at -80 °C and working stocks are stored for a few weeks at 5 mM aqueous solution at 4°C in opaque tubes given the photo-sensitive nature of this reagent. iN04 was purchased from ChemBridge Corporation (#7989904). Full-length recombinant human LRRK2[G2019S] was purchased from Invitrogen (#A15202). GZD-824 was purchased from Cayman Chemical (#21508) and calyculin A (#ab141784) was purchased from Abcam, Inc. Phorbol 12-myristate 13-acetate (PMA) (#P8139), interleukin-1A (#I2778), oligomycin (#75351), antimycin A (#A8674), AICAR (#A9978) and forskolin (#F6886) were all purchased from Sigma-Aldrich. Human EGF (#8916), and human IGF-1 (#3093) were purchased from cell signaling technology (CST). PKC-specific inhibitor GO6983 was purchased from Selleckchem (#S2911). ON-TARGETplus SMARTpool human LRRK1 (#L-005320-00-0005), human LRRK2 (#L-049666-00-0005) and non-targeting control pool (#D-001810-10-20) siRNA were purchased from Horizon Discovery.

### Plasmids

The following cDNA constructs were used for transfections: HA-empty vector (DU49303), HA-Rab10 (DU44250), HA-Rab8A (DU35414), HA-Rab7A wt (DU24784), HA-Rab7A[Q67L] (DU24813), HA-Rab29 (DU50222), GFP-Empty (DU44062), GFP-Rab7A wt (DU24785), GFP-Rab7A[S72A] (DU24814), GFP-LRRK2 wt (DU48062), GFP-LRRK2[D2017A] (DU48063), GFP-LRRK1 wt (DU30382), GFP-LRRK1[D1409A] (DU67084), GFP-LRRK1 [K746G] (DU67083), GFP-LRRK1[Y971F] (DU67289), GFP-LRRK1[K746G+Y971F] (DU66249), GFP-LRRK1[F1022C] (DU67082), GFP-LRRK1[G1411S] (DU67099), GFP-LRRK1[E929*] (DU66390), GFP-LRRK1[E929K] (DU66384), GFP-LRRK1[E929Q] (DU66391), GFP-LRRK1[Y971C] (DU66385), GFP-LRRK1[I1412F] (DU66395), GFP-LRRK1[E1980A-FS] (DU68320), FLAG-LRRK1[D1409A] (DU17400), FLAG-LRRK1[K746G] (DU67104), GFP-RILP (DU27496), HA-PPM1H wt (DU62789), HA-PPM1H[H153D] (DU62928), HA-PPM1J wt (DU68077), HA-PPM1J[H160D] (DU68170), HA-PPM1M wt (DU68124), and HA-PPM1M[H127D] (DU68165). These are available from MRC PPU Reagents and Services (https://mrcppureagents.dundee.ac.uk/). DNA constructs were amplified in Escherichia coli DH5α and purified using a Hi-Speed Plasmid Maxi Kit (Qiagen).

### Generation of Rabbit polyclonal pSer72 Rab7A

Rabbit immunization and rabbit antibody generation were performed by Abcam, Inc. (Burlingame, CA). The rabbits were immunized with peptides C-Ahx-AGQERFQS*LGVAFYR-amide and Ac-AGQERFQS*LGVAFYR-Ahx-C. Three subcutaneous injections were performed using the KLH immunogens, followed by two subcutaneous injections using the ovalbumin immunogens. At the time of each injection, an immunogen aliquot was thawed and combined with Freund’s Complete (initial immunisation) Adjuvant or Freund’s incomplete Adjuvant (for the subsequent injections). Serum bleeds were obtained after the fourth and fifth immunizations. Antibodies were affinity-purified using the phosphorylated epitopes as a positive selection and also passed through a column containing the dephosophorylated peptide as a negative selection step. The affinity-purified antibodies were evaluated in parallel with serum. The subsequent rabbit polyclonal antibodies were diluted in 5% (by mass) BSA (bovine serum albumin) in TBST Buffer (20 mM Tris pH 7.5, 0.15 M NaCl and 0.1% (by vol) Tween-20) and used for immunoblotting analysis at a final concentration of 1 μg/ml. The antibody was generated by The Michael J. Fox Foundation’s research tools program in partnership with Abcam. Development of a monoclonal antibody is underway. Please contact with questions.

### Antibodies

The sheep polyclonal antibodies for LRRK1 (sheep number S405C, 2^nd^ bleed), total Akt (sheep number S695B, 3^rd^ bleed), PINK1 (suitable for immunoblot) (sheep number S086D, 3^rd^ bleed), PINK1 (suitable for immunoprecipitation) (sheep number DA039, 3^rd^ bleed), Rab3A (sheep number SA098, 2^nd^ Bleed) and Rab43 (sheep number SA135, 6^th^ bleed) were generated by MRC Reagents and Services (https://mrcppureagents.dundee.ac.uk/). anti-LRRK1 was produced by immunization with C-terminal LRRK1 protein (residues 1695-end; GST tag cleaved) followed by 3 further injections 28 days apart, with bleeds performed seven days after each injection. Antibodies were affinity-purified from serum using LRRK1 antigen. For immunoblotting analysis, all sheep polyclonal antibodies were diluted in 5% (by mass) milk in TBST buffer. Rabbit monoclonal phospho-Ser935 LRRK2 (UDD2) was expressed and purified at University of Dundee as described previously [35]. The following antibodies were obtained from the indicated suppliers. Rabbit monoclonal MJFF-pRab10 (Thr73) (#ab230261, Abcam, Inc. [36]), rabbit monoclonal MJFF-pRab8 (Thr72) (#ab230260, Abcam, Inc. [36]), mouse monoclonal MJFF-total Rab10 antibody (#0680-100/Rab10-605B11, nanotools [36]), mouse monoclonal anti-LRRK2 (#75-253, NeuroMab), mouse monoclonal anti-Rab8A (#WH0004218M2, Sigma-Aldrich), mouse anti-α-Tubulin (#2144, Cell Signalling Technologies (CST)), rabbit polyclonal pERK1/2 (T202, Y204) (#9101, CST), rabbit monoclonal pEGFR (Y1173) (#4407, CST), rabbit monoclonal pAMPK (Thr172) (#4188, CST), mouse monoclonal total AMPK (CST, #2793), rabbit polyclonal pACC (Ser79) (CST, #3661), rabbit monoclonal pVASP (Ser157) (#84519, CST), rabbit polyclonal p-Ser PKC Substrate (#2261, CST), rabbit monoclonal pTBK1 (Ser172) (#5483, CST), rabbit polyclonal total TBK1 (#3013, CST), rabbit monoclonal pP105 (Ser933) (#4806, CST), rabbit monoclonal pP38 (Thr180, Tyr182) (#4631, CST), mouse monoclonal anti-Total Ub (#646302, BioLegend), mouse monoclonal anti-OPA-1 (#612607, BD Biosciences), mouse anti-GAPDH (#sc-32233, Santa Cruz Biotechnology), and rabbit anti-GFP (#PABG1, Chromotek), Rat anti-HA (#ROAHAHA, Sigma), mouse monoclonal anti-Rab7A (#R8779, Sigma), rabbit polyclonal anti-pAkt (Ser473) (#9271, CST), mouse monoclonal VPS35 (#SMC-605D, StressMarq Biosciences). Rabbit monoclonal anti-Total Rab35 (clone ID: 25) was kindly provided by Abcam, Inc. and will be available soon. Rabbit polyclonal pUbiquitin (Ser65) (21^st^ Century Biochemicals Inc.) was kindly provided by the Muqit lab and described previously [37]. All antibodies were diluted in 5% (by mass) BSA (bovine serum albumin) in TBST Buffer and used for immunoblotting analysis at a final concentration of 1 μg/ml.

### Mice

Heterozygous LRRK1 (wild type/ knock-out) mice were purchased from the JAX laboratory (https://www.jax.org/strain/016120). Mice were maintained under specific pathogen-free conditions at the University of Dundee (U.K.). All animal studies were ethically reviewed and carried out in accordance with Animals (Scientific Procedures) Act 1986 and regulations set by the University of Dundee and the U.K. Home Office. Animal studies and breeding were approved by the University of Dundee ethical committee and performed under a U.K. Home Office project license. Mice were multiply housed at an ambient temperature (20–24°C) and humidity (45–55%) and maintained on a 12 h light/12 h dark cycle, with free access to food (SDS RM No. 3 autoclavable) and water.

### LRRK1 KO Primary MEF generation

Wild-type, heterozygous, and homozygous LRRK1 knock-out mouse embryonic fibroblasts (MEFs) were isolated from littermate-matched mouse embryos at day E12.5 resulting from crosses between heterozygous wild-type/KO mice as described previously [38]. Genotyping of mice, embryos and MEFs was performed by PCR using genomic DNA, Primer 1 (12041 com: 5’ TCACTGGAAGGCAAGGACAGTTGG 3’), Primer 2 (12042 mut rev: 5’ CCAACACATCCAACCTTTTCTTTGC 3’), and Primer 3 (12043 wt rev: 5’ CAGGATGCTTCACAGACAGG 3’). The wild type sequence leads to a 368bp species whilst the knock-out produces a 220bp species.

### Cell Culture and Lysis

HEK293, HaCaT and HeLa cells were purchased from ATCC and cultured in DMEM containing 10% (by vol) fetal calf serum, 2 mM L-glutamine, 100 U/ml penicillin, and 100 μg/ml streptomycin. MEFs were grown in DMEM containing 10% (by vol) fetal calf serum, 2 mM L-glutamine, 100 U/ml penicillin, and 100 μg/ml streptomycin supplemented with non-essential amino acids and 1 mM sodium pyruvate. All cells were incubated at 37°C, 5% CO2 (by vol) in a humidified atmosphere and regularly tested for mycoplasma contamination. Cells were lysed in an ice-cold lysis buffer containing 50 mM Tris–HCl, pH 7.4, 1% Triton X-100 (by vol), 10% glycerol (by vol), 150 mM NaCl, 1 mM sodium orthovanadate, 50 mM NaF, 10 mM 2-glycerophosphate, 5 mM sodium pyrophosphate, 1 μg/ml microcystin-LR, and complete EDTA-free protease inhibitor cocktail (Roche). Lysates were clarified by centrifugation at 20,800 g at 4°C for 10 min and supernatant protein concentrations quantified using the Bradford assay.

### Transient overexpression experiments

24 h prior to transfection, cells were seeded into 6 well plates at 3 × 10^5^ cells/well. Cells were transfected using Polyethylenimine [39] employing 1.6 µg of LRRK1/LRRK2 plasmids, 0.4 µg of the Rab protein plasmid. For phosphatase experiments, cells were transfected with 1.0 µg of LRRK1 plasmids, 0.25 µg of Rab protein plasmid and 0.5 µg of phosphatase plasmids. 24 h post-transfection, cells were treated with the indicated concentrations of inhibitors dissolved in DMSO (0.1% (by vol) final concentration) for 2 h prior to cell lysis. An equivalent volume of DMSO was added to cells not treated with an inhibitor.

### siRNA knockdown experiments

HaCaT cells were seeded in 6-well plates at 2×10^5^ cells/well. After 24 h cells were either left untransfected or transfected using 7.5 µl Lipofectamine RNAi Max (Thermofisher, #13778150) and 10 nM of LRRK1, LRRK2 or non-targeting control siRNA per well. Cells were then cultured for a further 60 h. Cells were subsequently deprived of serum for 16 h and then treated in the presence or absence of 50 ng/ml EGF for 15 min and then lysed.

### HA-Rab7A and GFP-RILP Immunoprecipitation

Cells were transiently transfected as described above. 24 h post transfection, cells were harvested in lysis buffer and lysates clarified by centrifugation at 17,000 g for 10 min at 4 °C. 500 µg of whole cell lysate was incubated with 15 µl of packed beads of either nanobody anti-GFP binder-Sepharose or anti-HA frankenbody-Sepharose beads (generated by the MRC PPU Reagents and Services) for 1 h. Bound complexes were recovered by washing the beads three times with 50 mM Tris-HCl pH 7.5 150 mM NaCl before eluting with 2x SDS-PAGE sample buffer supplemented with 0.1% (by vol) 2-mercaptoethanol. The samples were denatured at 70°C for 10 min and the resin was separated from the sample by centrifugation through a 0.22 μm Spin-X® column (CLS8161, Sigma).

### PINK1 Immunoprecipitation

For immunoprecipitation of endogenous PINK1, primary MEFs were first treated either with 1 µM oligomycin and 10 µM antimycin A or with DMSO for 24 h. Subsequently, 500 µg of whole-cell lysate was incubated overnight at 4°C with 10 µg of PINK1 antibody (DA039, MRC PPU reagents and Services) coupled to Protein A/G beads (10 µl of beads per reaction, Amintra). The immunoprecipitants were washed three times with lysis buffer containing 150 mM NaCl and eluted by resuspending in 20 µl of 2× LDS sample buffer and incubating at 37°C for 15 min under constant shaking (2,000rpm) followed by the addition of 0.1% (by vol) 2-mercaptoethanol.

### Ubiquitin capture assay

For ubiquitin capture, primary MEFs were treated either with 1 µM Oligomycin and 10 µM Antimycin or with DMSO for 24 h and subsequently 500 µg of whole-cell lysate was used for pulldown with HALO-MultiDsk beads. In particular, MultiDsk (5x UBA domain of Dsk2 from yeast) was expressed as N-terminal GST and HALO tag recombinant protein in *E. coli* BL21 DE3, affinity purified on GSH –Sepharose. This was recovered by incubating the resin with TEV-protease for 3h and dialysed into 50 mM HEPES, 150 mM NaCl, 1 mM DTT). HALO-MultiDsk (2 mg) was incubated with 200 µl of HaloLink resin (Promega) in binding buffer (50 mM Tris–HCl pH 7.5, 150 mM NaCl, 0.05% (by vol) NP-40) overnight at 4 °C under agitation. HALO-TUBE beads (20 µl) were then added to whole-cell lysates from MEFs lysed as described above except lysis buffer supplemented with 100 mM chloroacetamide to inhibit cysteine-dependent deubiquitylating enzymes and incubated at 4 °C overnight under agitation. Beads were washed three times with lysis buffer containing 150 mM NaCl and eluted by resuspending in 20 µl 2X LDS sample buffer and incubating at 37 °C for 15 min under constant shaking (2,000rpm) followed by the addition of 0.1% (by vol) 2-mercaptoethanol.

### Immunoblotting

Lysates were incubated with a quarter of volume or 4X SDS-PAGE sample buffer [50 mM Tris–HCl, pH 6.8, 2% (by mass) SDS, 10% (by vol) glycerol, 0.02% (by mass) Bromophenol Blue and 1% (by vol) 2-mercaptoethanol] (Novex) and heated at 95 °C for 5 min. For normal SDS-PAGE, 10–20 μg samples were loaded on to 4-12% Bis-tris gradient gels and electrophoresed at 180 V for 90 min. Proteins were transferred onto nitrocellulose membranes at 90 V for 90 min on ice in transfer buffer [48 mM Tris/HCl, 39 mM glycine and 20% (by vol) methanol]. Transferred membranes were blocked with 5% (by mass) milk in TBST Buffer at room temperature for 60 min. Membranes were then incubated with primary antibodies diluted in blocking buffer overnight at 4 °C. After washing in TBST, membranes were incubated at room temperature for 1 h with near-infrared fluorescent IRDye antibodies (LI-COR) diluted 1:10000 in TBST and developed using the LI-COR Odyssey CLx Western Blot imaging system and signal quantified using the Image Studio software

### Phostag Immunoblot Analysis

Phos-tag SDS-PAGE was carried out as described previously [18]. Samples were supplemented with 10 mM MnCl_2_ before loading. Gels for Phos-tag SDS–PAGE consisted of a stacking gel [4% (by mass) acrylamide, 125 mM Tris–HCl, pH 6.8, 0.1% (by mass) SDS, 0.2% (by vol) N,N,N’,N’-tetramethylethylenediamine (TEMED) and 0.08% (by mass) ammonium persulfate (APS)] and a separating gel [12% (by mass) acrylamide, 375 mM Tris/HCl, pH 8.8, 0.1% (by mass) SDS, 75 mM Phos-tag acrylamide, 150 mM MnCl_2_, 0.1% (by vol) TEMED and 0.05% (by mass) APS]. 10– 30 μg of samples were loaded and electrophoresed at 70V for 30 min and at 120V for 2 h in running buffer [25 mM Tris/HCl, 192 mM glycine and 0.1% (by mass) SDS]. Gels were washed in transfer buffer containing 10 mM EDTA and 0.05% (by mass) SDS three times for 10 min, followed by one wash in transfer buffer containing 0.05% (by mass) SDS for 10 min. Proteins were transferred onto nitrocellulose membranes (Amersham Protran 0.45 μm NC; GE Healthcare) at 100 V for 180 min on ice in the transfer buffer without SDS/EDTA. Subsequent immunoblotting was performed as previously described.

### Immunofluorescence

For analysis of Rab29 co-localization with LRRK1 and LRRK2, HeLa cells were seeded in 6 well plates on glass coverslips (VWR, 631-0125; square 22×22 mm thickness 1.5). Cells were transfected using Polyethylenimine method [39] employing 1.6 μg of LRRK1/LRRK2 plasmids and 0.4 μg of either HA-Rab29 or HA-empty. 24 h post transfection cells were fixed with 4% (by vol) paraformaldehyde at room temperature for 10 min. Next, cells were permeabilized using 1% (by vol) NP-40 in PBS for 10 min followed by blocking with 1% (by mass) BSA in PBS. Slides were incubated with mouse monoclonal anti-HA (#ab18181, Abcam, Inc.) primary antibody, diluted 1:1000 in 1% (by mass) BSA in PBS buffer (137 mM NaCl, 2.7 mM KCl, 10 mM Na_2_HPO_4_, 1.8 KH_2_PO_4_, pH 7.4) for 1 h at room temperature followed with 3 times for 15 min washes with 0.2% (by mass) BSA in PBS (no antibody used for GFP, fluorescence of GFP was measured directly). Then slides were incubated with Alexa Fluor® 405 goat anti-mouse secondary antibody (Invitrogen A-31553) diluted 1:500 and HCS CellMask™ Deep Red Stain (Invitrogen, H32721) at 0.5 µg/mL dilution in 1% (by mass) BSA in PBS for 1 h. Slides were washed 3 times for 15 min with 0.2% (by mass) BSA in PBS and rinsed with distilled water just before mounting. VECTASHIELD® HardSet™ Antifade Mounting Medium (Vector Laboratories, H-1400) was used to mount the coverslips on glass slides (VWR, 631-0117). Slides were then imaged using Leica TCS SP8 MP Multiphoton Microscope using a 40x oil immersion lens choosing the optimal imaging resolution with 1-pixel size of 63.3 nm x 63.3 nm.

### Expression and purification of Rab proteins

4 × 500 ml of Lysogeny broth containing 100 μg/ml ampicillin antibiotic was inoculated with a single colony of BL21-CodonPlus(DE3)-RIPL strain of *E. coli* transformed with plasmids encoding for the expression of His-SUMO-Rab7A[wt] (Plasmid DU24781), His-SUMO-Rab7A[S72A] (DU24808) or His-SUMO-Rab10[wt] (Plasmid DU51062). Bacteria were cultured at 37 °C to an OD600 of 0.4-0.6, then temperature reduced to 15 °C and protein expression was induced by addition of Isopropyl β-D-1-thiogalactopyranoside at 250 μM. Cells were cultured for 16 h before harvesting by centrifugation at 4,200 x g for 20 mins at 4 °C. The pellet was resuspended in 200 ml ice cold *E. coli* lysis buffer [50 mM Tris/HCl pH7.5, 100 mM NaCl, 1 mM MgCl_2_, 10 µM GDP, 2 mM TCEP (tris(2-carboxyethyl)phosphine)]. Cells were lysed using a cell disruptor (passing sample through twice) and extracts clarified by centrifugation at 15, 000 rpm for 20 mins at 4 °C. Subsequent supernatants were run on 15 ml of Cobalt-Agarose resin (Amintra Cobalt NTA Affinity Resin, Expedeon), previously equilibrated in *E. coli* lysis buffer, at 4 °C for 90 min. The column was then washed with 20 column volumes of High Salt Wash Buffer [50 mM Tris/ HCl pH7.5, 500 mM NaCl, 1 mM MgCl_2_, 10 µM GDP, 2 mM TCEP] until no unbound protein was present in the flow-through. The column was then washed with 5 column volumes of Low Salt Wash buffer [50 mM Tris/ HCl pH7.5, 150 mM NaCl, 2 mM MnCl_2_, 10 µM GDP), 2 mM TCEP]. His-SENP1 (DU39129) (1:30 dilution) was then added to the column overnight at 4 °C to cleave the His-SUMO tag. Cleaved protein was collected and subjected to gel filtration on a Superdex 75 Hi load column, previously equilibrated in Equilibration buffer [50 mM Tris/HCl pH7.5, 150 mM NaCl, 2 mM MnCl2, 1 mM GTP, 0.5 mM TCEP]. 0.2 ml fractions were collected at a flow rate of 0.2 ml/min, with protein-containing fractions pooled, subjected to spin concentration using an Amicon Ultra 15 10 KDa and finally snap frozen.

### Cloning, expression and purification of LRRK1

The DNA coding for the human LRRK1 residues 28 to 2015 (OHu72031 from Genscript) was PCR-amplified using the forward primer TACTTCCAATCCATGGAGACGCTTAACGGTGCCGGGGAC and the reverse primer TATCCACCTTTACTGCTTTACCTTCTCTTGCGAGTGCAAGC. The T4 polymerase-treated amplicon was inserted into the transfer vector pFB-6HZB (SGC) by ligation-independent cloning. The resulting plasmid was utilized for the generation of recombinant Baculoviruses according to the Bac-to-Bac expression system protocol (Invitrogen). Exponentially growing Sf9 cells (2E06 cells/mL in Lonza Insect-XPRESS medium) were infected with high-titre Baculovirus suspension. After 66 h of incubation (27°C and 90 rpm), cells were harvested by centrifugation. The expressed protein construct contained an N-terminal His_6_-Z-tag, cleavable with TEV protease. For LRRK1 purification, the pelleted Sf9 cells were washed with PBS, re-suspended in lysis buffer (50 mM HEPES pH 7.4, 500 mM NaCl, 20 mM imidazole, 0.5 mM TCEP, 5% (by vol) glycerol) and lysed by sonication. The lysate was cleared by centrifugation and loaded onto a Ni NTA column. After vigorous rinsing with lysis buffer the His_6_-Z-tagged protein was eluted in lysis buffer containing 300 mM imidazole. Immediately thereafter, the eluate was diluted with buffer containing no NaCl, to reduce NaCl to 250 mM and loaded onto an SP-Sepharose column. His_6_-Z-TEV-LRRK1 was eluted with a 250 mM to 2.5 M NaCl gradient and treated with TEV protease overnight to cleave the His_6_-Z-tag. Contaminating proteins, the cleaved tag, uncleaved protein and TEV protease were removed by another combined SP-Sepharose Ni NTA step. Finally, LRRK1 was concentrated and subjected to gel filtration in storage buffer (20 mM HEPES pH 7.4, 150 mM NaCl, 0.5 mM TCEP, 5% glycerol) using an AKTA Xpress system combined with an S200 gel filtration column. The final yield as calculated from UV absorbance was 0.1 mg/L.

### LRRK1 and LRRK2 in vitro kinase assays

Kinase assays were set in a 20 μl final reaction mixture containing purified Rab proteins (2 μM, 1 μg), recombinant LRRK1 (20 nM, 100 ng, residues 28-2015) or full-length LRRK2[G2019S] (20 nM, 100 ng) and 50 mM Tris–HCl pH 7.5, 0.1 mM EGTA, 10 mM MgCl_2_ and 1 mM ATP. The kinase reactions were carried out at 30 °C for the indicated times and reactions terminated by the addition of 4x SDS– PAGE sample buffer containing 1% (by vol) 2-mercaptoethanol. 20% of total reaction vol was used for Phos-tag Coomassie staining analysis; <10% was used for conventional immunoblot analysis.

### Phosphosite identification by MS and Edman sequencing

The phosphorylation sites on Rab7A were identified as previously described [16]. Purified Rab7A (12.5 μM, 7.5 μg) was phosphorylated using recombinant full-length wild type LRRK1 (80 nM, 400 ng) in a buffer containing 50 mM Tris-HCl pH 7.5, 10 mM MgOAc, 0.1 mM [γ-^32^P]ATP (specific activity: ∼3000 Ci/pmol) for 1 h at 30°C. The reactions were stopped by the addition of 4X SDS-PAGE sample buffer containing 1% (by vol) 2-mercaptoethanol, and samples were subjected to electrophoresis on a 4-12% SDS-PAGE gels. The gels were was stained with Coomassie blue and the band corresponding to Rab7A was excised and washed with water (10 min) and subsequently subjected to two rounds of washes with 100% acetonitrile and 50 mM Tris–HCl pH 8.0 (5 min per wash). Gel pieces were then reduced by incubation in 5 mM DTT in 50 mM Tris–HCl pH 8.0 at 65°C for 20 min. This was followed by alkylation of the samples by addition of 20 mM iodoacetamide in 50 mM Tris–HCl pH 8.0 for 20 min. The gel pieces were then washed with 100% acetonitrile, 50 mM Triethylammonium bicarbonate and again 100% acetonitrile (5 min each). Any remaining acetonitrile was removed with a pipette before adding trypsin (5 mg/ml in 50 mM Triethylammonium bicarbonate in a volume sufficient to cover the gel pieces) for overnight digestion at 30°C. The resulting tryptic peptides were extracted from the gel pieces by addition of 100% acetonitrile. These were subsequently resuspended in 0.1% (by vol) trifluoroacetate (TFA) in 5% (by vol) acetonitrile before being separated on a reverse-phase HPLC Vydac C18 column (Separations Group) with an on-line radioactivity detector. The column was equilibrated in 0.1% (v/v) trifluoroacetic acid and developed with a linear acetonitrile gradient at a flow rate of 0.2 ml/min. Fractions (0.1 ml each) were collected and analyzed for ^32^P radioactivity by Cerenkov counting with a tricarb scintillation counter. All the identified phosphopeptide fractions were analyzed by liquid chromatography (LC)-MS/MS using a Thermo U3000 RSLC nano liquid chromatography system (Thermo Fisher Scientific) coupled to a Thermo LTQ-Orbitrap Velos mass spectrometer (Thermo Fisher Scientific) to determine the primary sequence of the phosphopeptides. Data files were searched using Mascot (www.matrixscience.com) run on an in-house system against a database containing the appropriate Rab sequences, with a 10 ppm mass accuracy for precursor ions, a 0.6 Da tolerance for fragment ions, and allowing for Phospho (ST), Phospho (Y), Oxidation (M), and Dioxidation (M) as variable modifications. Individual MS/MS spectra were inspected using Xcalibur 2.2 (Thermo Fisher Scientific), and Proteome Discoverer with phosphoRS 3.1 (Thermo Fisher Scientific) was used to assist with phosphosite assignment. The site of phosphorylation of ^32^P-labeled peptides was determined by solid-phase Edman degradation on a Shimadzu PPSQ33A Sequencer of the peptide coupled to Sequelon-AA membrane (Applied Biosystems) as described previously [40].

### In-gel digestion for LC/MS-MS

In-gel digestion and targeted mass spectrometry was performed as described [41]. Briefly, 50 µg of lysates from each of the two independent clones of LRRK1 KO and LRRK1 wt cells, as technical duplicates were reduced by adding 5 mM Dithiothreitol and incubated at 56 °C for 20 min on a Thermo mixer with continuous agitation at 800 rpm. The tubes were brought to room temperature and alkylated by adding 20 mM Iodoacetamide at room temperature in the dark on a Thermomixer with continuous agitation for 20 min. The protein lysates were then resolved by SDS-PAGE. The gel was stained with colloidal Coomassie (NUPAGE), destained overnight and subjected to in-gel digestion. The region between 30 to 20 kDa was excised and destained with 200 µl, 40 mM Ammonium bicarbonate in 40% Acetonitrile buffer and incubated at room temperature on a Thermomixer with continuous agitation at 1200 rpm for 30 min. The gel pieces were completely destained by repeating the above step, then rehydrated by adding 200 µl of Acetonitrile and incubated on a Thermomixer. Acetonitrile was then removed, and slices subjected to vacuum drying for 10 min to ensure the complete Acetonitrile removal. The trypsin was prepared in 0.5% (by mass) sodium deoxycholate (final) in 20 mM TEABC buffer, 500 ng trypsin to each sample was added and kept under incubation at 37°C overnight on a Thermomixer with a continuous agitation at 1200 rpm. The tryptic peptides were extracted by adding 200 µl of extraction buffer (99% Isopropanol in 1% (by vol) TFA, incubated on a Thermomixer with an agitation at 1200 rpm for 20 min. The extraction was completed by repeating again and the eluate was directly loaded on in-house prepared SDB-RP Stage-tips and subjected to the SDC-RP clean up. Two disks were prepared by punching with 16-gauge needle and loaded on to 250 µl pipette tips. The peptide digest was loaded and centrifuged at room temperature at 2,000g for 10 min. The flow through was reapplied and subsequently the stage tips were washed once with 150 µl wash buffer I (99% Isopropanol in 1% TFA) and washed again with 150 µl wash buffer II (3% Acetonitrile in 0.2% TFA). All the centrifugation was done at room temperature for 10 min. The peptides were eluted by adding 60 µl of elution buffer I (50% Acetonitrile in 1.25% Ammonium Hydroxide), further, the elution was repeated by adding 60 µl of elution buffer II (80% Acetonitrile in 1.25% Ammonium Hydroxide). All the centrifugation was done at room temperature at 1500g for 4 min. The peptide eluate was immediately snap frozen and vacuum dried for 90 min.

### LC-MS/MS and Data analysis

We undertook a multiplexed parallel reaction monitoring approach to identify and quantify Rab proteins phosphorylated on their effector binding motifs in wild type and LRRK1 knock-out MEFs. We synthesized heavy labeled stable isotope synthetic phosphopeptides encompassing the tryptic peptide sequences of pRab3, pRab7A, pRab8, pRab10, pRab35 and pRab43 (>98% purity, AAA analyzed from JPT Peptide Technologies, GmbH, Germany, sequences and masses listed on Supplementary File 1). 25 fmol the phospho Rab peptide mixture was spiked into the in-gel digested samples prior to loading into Evotips as described [42]. The Evotips (evosep.com, Odense, Denmak) were used according to the manufacturer. The disposable Evotips were loaded onto the EvoSep liquid chromatography (LC) system. EvoSep LC system employs a preformed gradient to ensure the focused separation of the peptides into the mass spectrometer and to minimize sample to sample carryover and overhead time between runs. The instrument was run using the “60 sample per day, 21 min run” method. Peptides were separated on 8 cm analytical columns (EV-1064, Dr Maisch C18 AQ, 3um beads, 100um ID, 8cm long). The EvoSep LC system was interfaced with QE-HFX mass spectrometer (Thermo Fisher Scientific, Bremen). The inclusion list for heavy and endogenous phosphopeptides along with the scheduled retention times for the targeted PRM method was generated using Skyline software imported directly into the QE-HFX Excalibur method. A full scan, MS1 between 300 to 800 m/z mass range was acquired at 120,000 at 200 m/z resolution and a PRM scan was acquired at 30,000 at 200 m/z resolution. The quadrupole isolation width of 0.7 m/z was set, and the peptides were fragmented using 27% normalized higher energy collisional dissociation (HCD) by maintaining the loop count to 10 (which acquires 10 MS2 scans per duty cycle). Both the AGC target and Ion injection times were set as 3E6 for MS1 and 1E5 for PRM and 50 ms for MS1 and 250 ms for PRM scan. The mass spectrometry raw data were imported into Skyline version 19.1.0.193 (University of Washington, Seattle USA,) software suite [43]. Skyline automatically processes and generates the peak area values for both heavy internal standards and endogenous phosphopeptides by selecting most intense fragment ions based on the pre-generated library which is a mix of heavy phosphopeptides. The elution profiles of each phosphopeptide was further manually verified and peak area values of both internal standard and endogenous phosphopeptides were used to generate the ratio values for each phosphopeptide. These ratio values were then compared between samples derived from the wild type and LRRK1 knock-out MEFs, enabling the relative levels of pRab3 (Thr86 in Rab3A,B,D & Thr94 in Rab3C), pRab7A (Ser72), pRab8 (Thr72), pRab10 (Thr73), pRab35 (Thr72) and pRab43 (Thr82) to be quantified.

## Results

### Ser72 phosphorylation of endogenous Rab7A is decreased in LRRK1 knock-out MEFs

To test whether LRRK1 might regulate the phosphorylation of Rab proteins, we generated littermate-matched wild type, heterozygous and LRRK1 homozygous knock-out mouse embryonic fibroblasts (MEFs), derived from previously described heterozygous LRRK1 knockout mice purchased from JAX laboratory (https://www.jax.org/strain/016120). To measure phosphorylation of endogenous Rab proteins in these cells, we used a combination of previously described quantitative targeted multiplex parallel reaction monitoring (PRM) mass spectrometry assays [15, 41]. For this experiment, we analyzed 2 independent clones for each genotype, derived from separate embryos. Briefly, 50 μg of each extract was electrophoresed on an SDS-polyacrylamide gel and the region encompassing Rab proteins (20-30 kDa) was excised and digested with trypsin. A mixture of heavy labelled phosphorylated peptides that encompass the tryptic peptide of Rab3, Rab7A, Rab8, Rab10, Rab35 and Rab43 (25 fmol of each) was spiked in. The resulting sample was subjected to targeted LC-MS/MS analysis enabling the levels of basal endogenous phosphorylation of each Rab protein in wild type and LRRK1 knock-out cells to be quantified. The analysis revealed that in each of the independent clones there was a ∼4-fold decrease in the phosphopeptide encompassing Rab7A phosphorylated at Ser72 in the LRRK1 knock-out compared to wild type cells (Fig 1A, upper panel). Ser72 is the residue equivalent to the LRRK2 phosphorylation sites on Rab8A and Rab10. The levels of peptides corresponding to Rab3, Rab8, Rab10, Rab35 and Rab43 phosphorylated residues were similar in wild type and LRRK1 knock-out extracts. Immunoblot analysis revealed that the levels of total Rab7A, Rab8A, Rab10, Rab35 and Rab43 were also similar in the wild type and knock-out MEFs, though total Rab3A levels were moderately lower in one of the knock-out clones (Fig 1A, lower panel).

**Figure 1.**
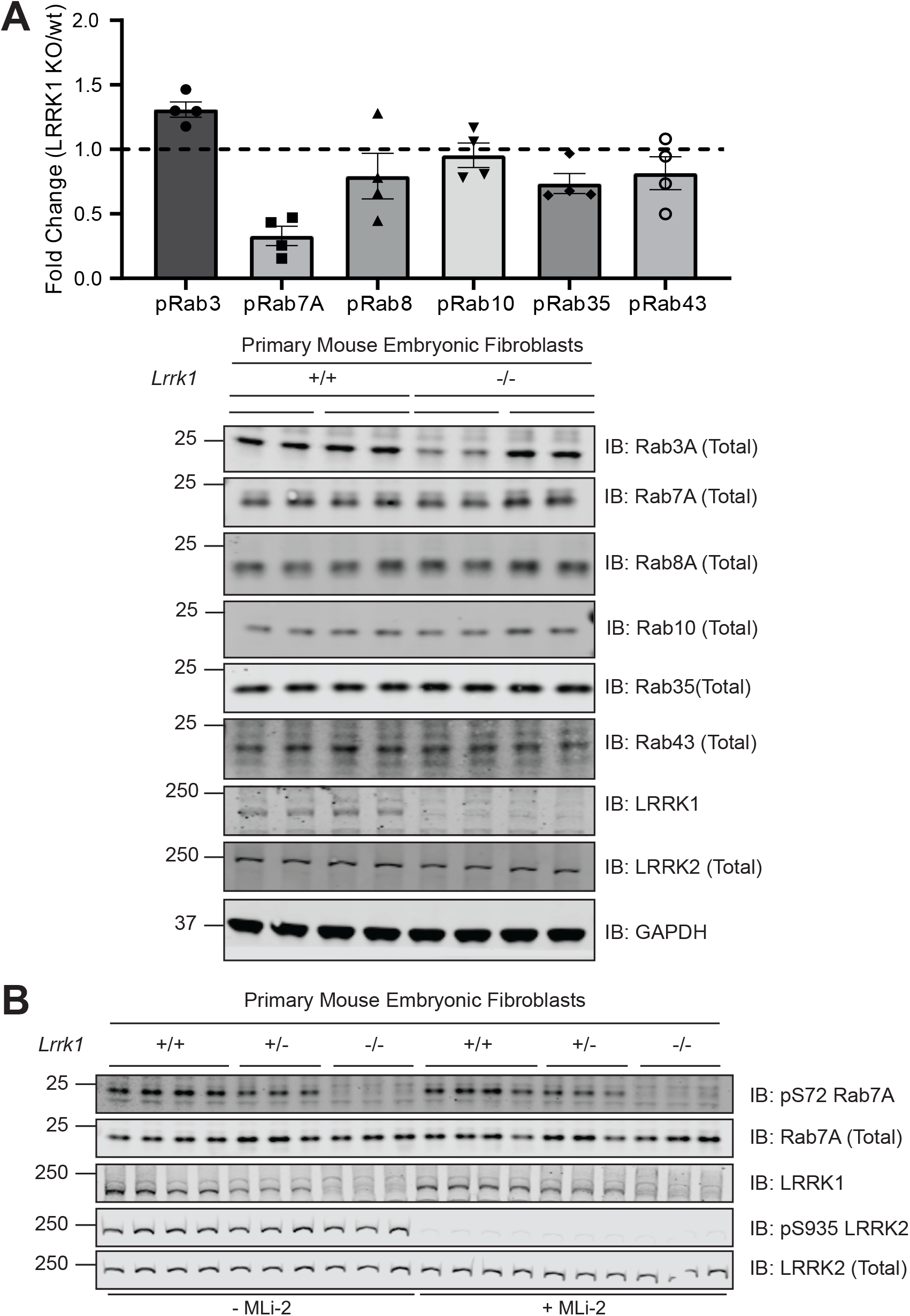
LRRK1 phosphorylates endogenous Rab7A at Ser72. Wild type and homozygous LRRK1 knock-out primary MEFs cultured in 10% (by vol) serum were lysed. (1A, upper panel) 50 µg of extracts from two independent clones were subjected to SDS-polyacrylamide gel electrophoresis and the region encompassing Rab proteins (20-30 kDa) was excised and subjected to in-gel digestion using trypsin. The extracted peptides were spiked with 25 femto moles of heavy phosphorylated Rab3, Rab7a, Rab8, Rab10, Rab35 and Rab43. Samples were analyzed using Parallel reaction monitoring (PRM) acquisition mode on a QE-HFX mass spectrometer. The raw data was processed using Skyline software and the relative expression of each pRab was determined between LRRK1 KO and wt. n=4, Each biological replicate analyzed as technical duplicate of each sample and the individual data marked with a black circle and the data presented as mean ± SEM. (1A, lower panel) The MEF extracts (20 μg) from 3 independent clones for wild type and homozygous knock-out clones were subjected to immunoblot analysis with the indicated antibodies (all at 1 µg/ml). Each lane represents cell extract obtained from a different clone. The membranes were developed using the LI-COR Odyssey CLx Western Blot imaging system. (B) As in A, except that 3 independent clones of littermate-matched wild type, heterozygous LRRK1 knock-out (+/-) and homozygous knock-out (-/-) primary MEFs were treated ± 200 nM MLi-2 for 30 min.

To further demonstrate that basal phosphorylation of Rab7A at Ser72 was reduced in LRRK1 knock-out cells, we utilized an affinity purified, phospho-specific pRab7A (Ser72) rabbit polyclonal antibody that we generated (see Materials and Methods). Using this antibody, we were able to detect endogenous Rab7A phosphorylation at Ser72 in all wild type MEF clones, that was virtually abolished in all of the LRRK1 knock-out cells (Fig 1B). We also studied LRRK1 heterozygous MEFs which displayed ∼50% lower levels of total LRRK1 and phosphorylated, compared to the wild type cells (Fig 1B). We observed that the LRRK1 deficient cells expressed similar levels of LRRK2 protein that was also similarly phosphorylated at Ser935. Treatment of MEFs with the LRRK2 MLi-2 inhibitor [44], did not impact Rab7A phosphorylation at Ser72, but consistent with previous work [45], inhibited phosphorylation of LRRK2 at Ser935 (Fig 1B).

### Overexpression of LRRK1 induces phosphorylation of Rab7A but not Rab8A or Rab10

We next explored whether LRRK1 overexpression could induce phosphorylation of Rab7A at Ser72. We co-expressed wild type or kinase inactive LRRK1[D1409A] mutant with Rab7A in HEK293 cells. Immunoblot analysis of cell extracts revealed that wild type LRRK1 but not kinase inactive LRRK1[D1409A] induced significant phosphorylation of Rab7A (Fig 2A). In parallel experiments wild type LRRK2 that was expressed at a similar level to LRRK1 did not induce phosphorylation of Rab7A (Fig 2A). We also demonstrated that wild type LRRK1 does not significantly phosphorylate Rab8A at Thr72 or Rab10 at Thr73 when overexpressed in HEK293 cells under conditions which LRRK2 efficiently phosphorylates these substrates (Fig 2A). To confirm the specificity of the pRab7A(Ser72) phospho-specific antibody, we mutated Ser72 to Ala in Rab7A, and demonstrated that no phospho-signal was detected when this protein was co-expressed with wild type LRRK1 (Fig 2B).

**Figure 2.**
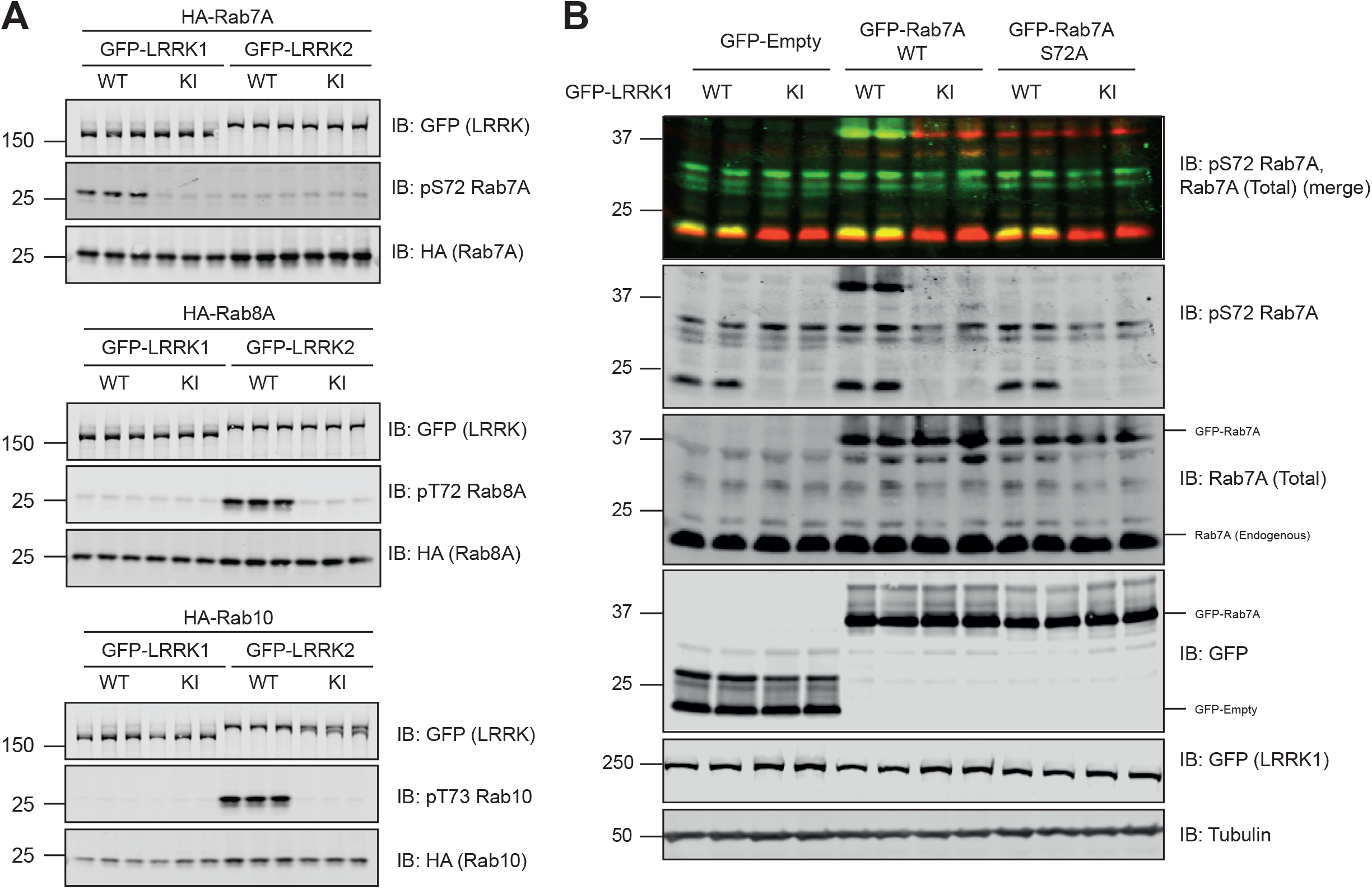
Further evidence that LRRK1 specifically phosphorylates Rab7A. (A & B) HEK293 cells were transiently transfected with the indicated plasmids encoding for wild type (wt) or kinase inactive (KI, D1409A) LRRK1 or LRRK2 and the indicated wild type or mutant Rab protein. 24 h post-transfections the cells were lysed and extracts (20 μg) from a triplicate experiment in which cells cultured in separate dishes were subjected to immunoblot analysis with the indicated antibodies (all at 1 µg/ml). Each lane represents cell extract obtained from a different replicate. The membranes were developed using the LI-COR Odyssey CLx Western Blot imaging system.

### LRRK1 directly phosphorylates Rab7A at Ser72 in vitro

To test whether LRRK1 can phosphorylate Rab7A directly, we first expressed and purified to homogeneity recombinant LRRK1 (lacking the first 27 residues i.e. residues 28 to 2015/end) in Sf9 insect cells (SFig 1). We also purified recombinant wild type Rab7A and Rab7A[S72A] mutant. Immunoblot analysis revealed that LRRK1 (20 nM) in the presence of MgATP, readily phosphorylated wild type Rab7A complexed to GDP (2 μM) (Fig 3A). To probe stoichiometry of Rab7A phosphorylation by LRRK1, we utilized the “phos-tag” approach that has previously been exploited to resolve dephosphorylated and LRRK2 phosphorylated Rab proteins by electrophoresis [18]. This revealed that a significant proportion of the wild type Rab7A was phosphorylated by LRRK1 under the kinase assay conditions we deployed (Fig 3A). In parallel experiments, immunoblot and phos-tag analysis revealed that the Rab7A[S72A] mutant was not phosphorylated by LRRK1 (Fig 3A). We also phosphorylated wild type Rab7A with LRRK1 in the presence of *γ*^32^P-ATP (Fig 3B) and subjected the ^32^P-labelled Rab7A protein to classical trypsin digestion-phosphopeptide mapping analysis followed by Edman degradation sequencing and mass spectrometry analysis. This confirmed unambiguously that Rab7A was phosphorylated by LRRK1 at a single site, namely Ser72 (Fig 3C). We also found that LRRK1 does not phosphorylate Rab10 in vitro, and vice versa, LRRK2 does not phosphorylate Rab7A (Fig 3D). These results are consistent with a previous report that LRRK1 phosphorylates Rab7A at Ser72 in vitro [28].

**Figure 3:**
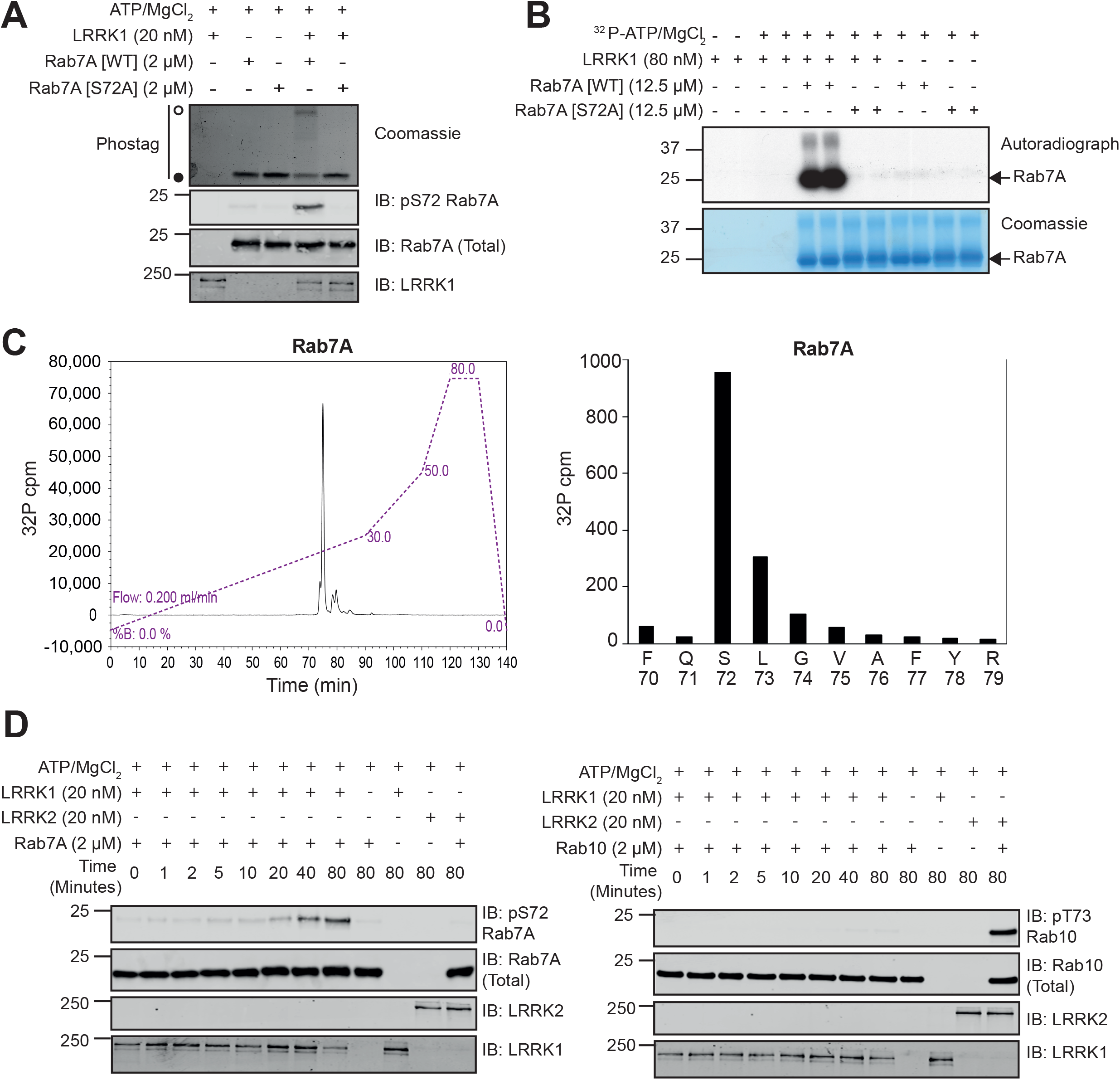
Recombinant LRRK1 phosphorylates Rab7A at Ser-72 in vitro. (A) Wild type (wt) Rab7A or Rab7A[S72A] was incubated in a phosphorylation reaction with recombinant LRRK1 as described in the materials and methods. Reactions were terminated after 60 min with SDS-Sample buffer. Samples subjected phos-tag electrophoresis followed by Coomassie staining (upper panel) or conventional immunoblot analysis with the indicated antibodies (all at 1 µg/ml) (lower panel). The membranes were developed using the LI-COR Odyssey CLx Western Blot imaging system. A Coomassie gel of the recombinant LRRK1 utilized in this experiment is shown in SFig 1. (B) As in (A) except that [*γ*^32^P]-ATP was employed for the phosphorylation reaction. Samples were resolved by SDS-PAGE and stained using Coomassie blue (upper band) and subjected to autoradiography (lower band). (C) Bands corresponding to ^32^P-Rab7A were excised from the gel and subjected to trypsin digestion. Resulting peptides were separated by reverse-phase C_18_chromatography and ^32^P radioactivity was detected using an online radioactivity detector. (D) The major peak of ^32^P-radioactivity was analyzed by Edman degradation as well mass spectrometry. Release of ^32^P radioactivity after each cycle of Edman degradation was determined and the amino acid sequence of the peptide determined by mass spectrometry analysis is shown. (E) Wild type Rab10 (upper panel) or Rab7A (lower panel) were incubated in a phosphorylation reaction with either recombinant LRRK1 or LRRK2 described in the materials and methods. Reactions were terminated at the indicated time-points with SDS-Sample buffer and analyzed by immunoblot analysis with the indicated antibodies (all at 1 µg/ml).

### Phorbol ester enhances LRRK1-mediated Rab7A phosphorylation in MEFs

As previous work had demonstrated that LRRK1 co-immunoprecipitates with components of the EGF receptor signalling pathway [24-26], we tested a range of agonists including EGF, IGF1, and phorbol esters to see whether any of these promote Rab7A phosphorylation in wild type MEFs. For each agonist we verified that it induced phosphorylation of a known component of its respective signalling pathway (SFig 2). These experiments revealed that the phorbol ester, phorbol 12-myristate 13-acetate (PMA, 30 min, 100 ng/ml) markedly stimulated Rab7A phosphorylation (Fig 4A, SFig 2). In contrast, none of the other agonists that we tested in MEFs increased Rab7A phosphorylation other than the potent PP1 and PP2A phosphatase inhibitor, calyculin-A (Fig 4A). In the LRRK1 knock-out cell line, PMA stimulation failed to induce Rab7A phosphorylation (Fig 4B); reduced phosphorylation of Rab7A was seen in heterozygous cells (Fig 4B). Time course analysis revealed that PMA induces Rab7A phosphorylation after ∼5 min, with maximal phosphorylation at 40 min that then declines to lower levels by 5 h (Fig 4C). We next pre-treated wild type MEFs with a panel of well characterized kinase inhibitors (Fig 4D, SFig 3). Only Go-6983 [46], a general PKC inhibitor, suppressed Rab7A phosphorylation induced by PMA (Fig 4D, SFig 3). We also studied Rab7A phosphorylation in human immortalized skin keratinocyte HaCaT cell line. EGF stimulation, but not phorbol ester, enhanced Rab7A phosphorylation (SFig 4A) and siRNA knock-down of LRRK1 inhibited Rab7A phosphorylation induced by EGF (SFig 4B).

**Figure 4.**
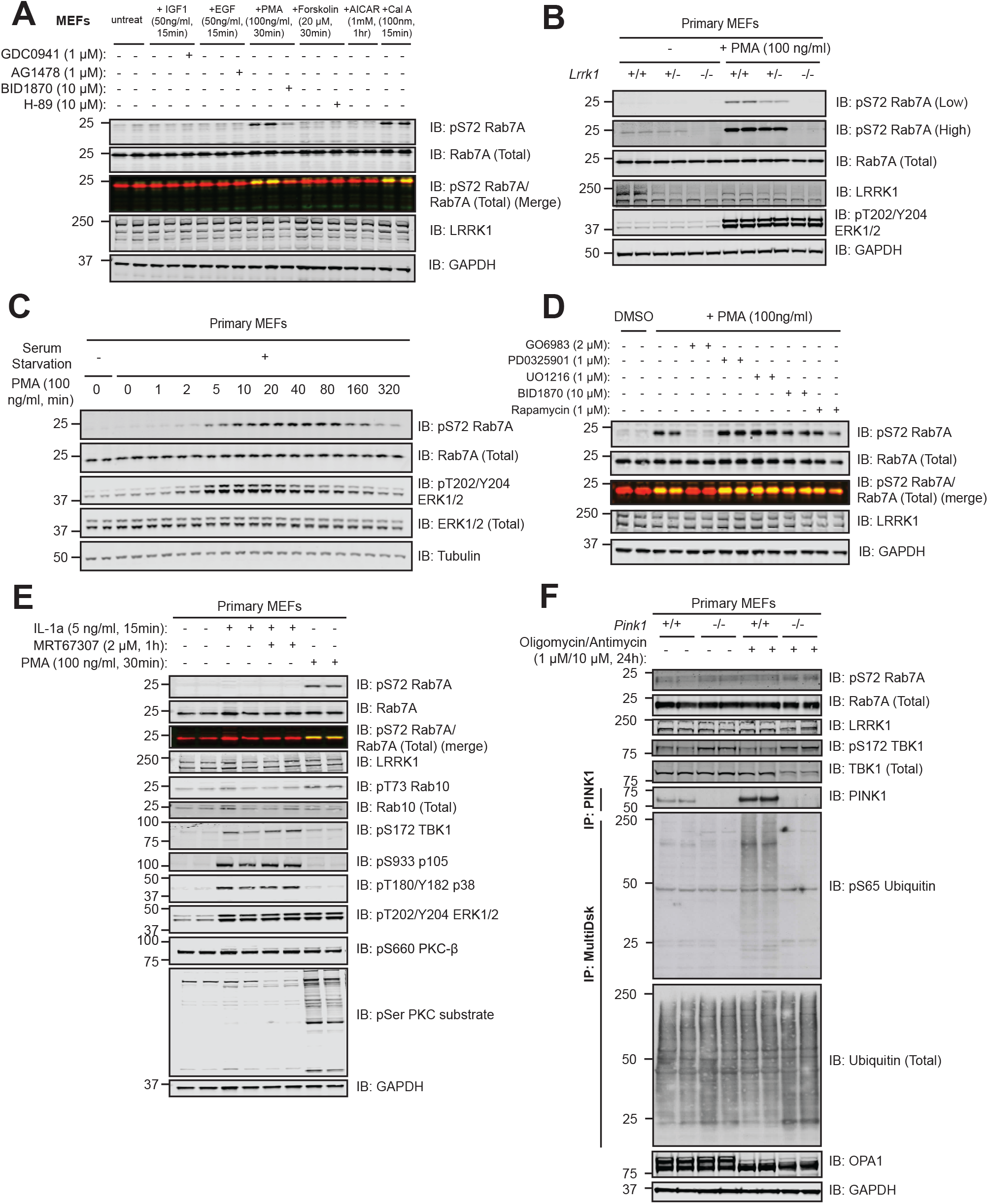
PMA stimulates LRRK1-dependent Rab7A phosphorylation. (A) Wild type MEFs were deprived of serum for 16 h and incubated for 1 h in the presence or absence of the dose of indicated inhibitor. Cells were next stimulated with the dose of indicated agonists for the times shown. Control blots for all agonists are included in SFig 2. Extracts (20 μg) from duplicate independent clones were subjected to immunoblot analysis with the indicated antibodies (all at 1 µg/ml). Each lane represents cell extract obtained from a different cell clone. The membranes were developed using the LI-COR Odyssey CLx Western Blot imaging system. (B) Wild type, heterozygous (+/-) and homozygous LRRK1 knock-out (-/-) primary MEFs were deprived of serum for 16 h and incubated in the presence or absence of 100 ng/ml PMA for 30 min and immunoblotted as in (A). (C) As in (A) except that wild type MEFs were treated with 100 ng/ml PMA for the indicated time-points. (D) As in (A), except that wild type MEFs were deprived of serum for 16 h and then incubated for 1 h in the presence or absence of the dose of indicated inhibitor. Cells were then stimulated with PMA (100 ng/ml) for 30 min. Control blots for all inhibitors are shown in SFig 3. (E) As in (A) except that cells were treated in the presence or absence of 2 μM MRT67307 for 1h and then stimulated with either 5 ng/ml interleukin-1a (IL-1A) for 15 min or 100 ng/ml PMA for 30 min. (F) As in (A) except that the previously described [51] wild type and homozygous PINK1 knock-out MEFs were treated in the presence or absence of 1 μM oligomycin and 10 μM antimycin A for 24h. For measurement of PINK1 stabilization, PINK1 was immunoprecipitated from extracts using PINK1 antibody (DA039, MRC PPU reagents and Services) coupled to Protein A/G beads (see materials and methods). For measurement of phosphorylation of ubiquitin at Ser65, the ubiquitin in the extracts was enriched by performing HALO-MultiDsk affinity purification (see materials and methods).

### PINK1 or TBK1 activation does not promote Rab7A phosphorylation in MEFs

Previous work had suggested that a PINK1-TBK1 signalling pathway was responsible for Rab7A phosphorylation in vivo [47, 48]. To investigate whether TBK1 induces phosphorylation of Rab7A in MEFs, we stimulated cells with interleukin-1A (IL-1A) a known activator of TBK1 in MEFs [49]. We confirmed that IL-1A stimulation (15 min, 5 ng/ml) promoted TBK1 activation as judged by an increase in Ser172 phosphorylation (Fig 4E), an autophosphorylation site that is used as a reporter for TBK1 activation [50]. However, Rab7A was not significantly phosphorylated following IL-1A treatment (Fig 4E). Moreover, PMA did not promote TBK1 phosphorylation at Ser172 (Fig 4E). We also stimulated MEFs with IL-1A for up to 5 h but no significant phosphorylation of Rab7A was observed at any timepoint studied, although it could be argued that the slightest increase beyond background levels was observed at the 20 min timepoint (SFig 5). We also treated previously reported wild type and PINK1 knock-out MEFs [51] with oligomycin (1 μM) and antimycin (10 μM) for 24 h to induce mitochondrial depolarization and activate the PINK1 protein kinase [37, 52]. PINK1 kinase was stabilized and activated in wild type but not PINK1 knock-out MEFs as judged by monitoring phosphorylation of ubiquitin at Ser65 (Fig 4F). Oligomycin and antimycin also induced cleavage of OPA-1, confirming mitochondrial depolarization (Fig 4F). However, oligomycin and antimycin A failed to induce phosphorylation of TBK1 or Rab7A in these experiments (Fig 4F).

### Identification of mutations that enhance LRRK1 activity

We next studied whether mutations equivalent to the common Parkinson’s causing LRRK2 mutations would affect LRRK1’s ability to phosphorylate Rab7A in cells. The mutants generated were LRRK1[K746G] (equivalent to LRRK2[R1441G]), LRRK1[F1022C] (equivalent to LRRK2[Y1699C]), LRRK1[G1411S] (equivalent to LRRK2[G2019S]) and LRRK1[I1412T] (equivalent to LRRK2[I2020T]) (Fig 5A, 5B). In addition, we also generated the previously described LRRK1[Y971F] mutant located within the COR domain (Fig 5B), reported to activate LRRK1 by blocking an inhibitory EGF receptor-mediated phosphorylation ([27, 28]). In overexpression studies, we observed that the LRRK1[K746G] and LRRK1[Y971F] mutations markedly increased Rab7A phosphorylation ∼5-fold (Fig 5C). The LRRK1[I1412T] mutation enhanced Rab7A phosphorylation more moderately (∼2-fold; Fig 5C). In contrast, the LRRK1[F1022C] (equivalent to LRRK2[Y1699C]) and LRRK1[G1411S] (equivalent to LRRK2[G2019S]) displayed similar activity to wild type LRRK1 (Fig 5C). We also generated a LRRK1[K746G+Y971F] double mutant but found that this displayed similar elevated activity as the single mutants (SFig 6).

**Figure 5.**
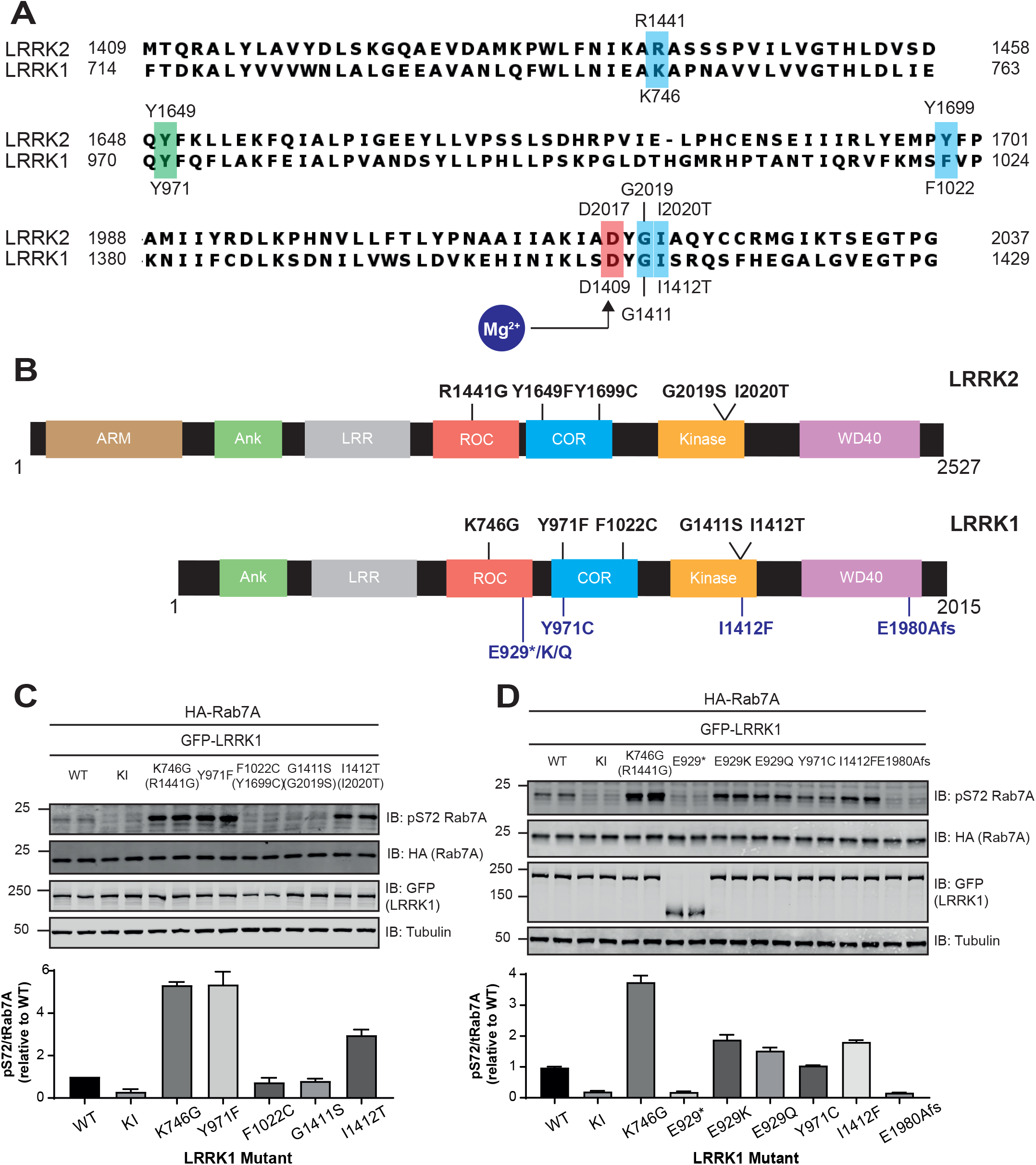
Identification of mutations that enhance LRRK1 activity. (A) Sequence alignment of LRRK2 and LRRK1. Residues corresponding to Parkinson’s mutations in LRRK2 are highlighted in blue. Tyr971 was reported to be an inhibitory phosphorylation site targeted by EGF receptor is indicated in green. (B) Domain arrangement of LRRK2 and LRRK1, with pathogenic and corresponding residues highlighted in black. Reported variants of LRRK1 are indicated in blue. (C&D) HEK293 cells were transiently transfected with the indicated plasmids encoding for wild type (wt) and indicated mutant of LRRK1 and wild type Rab7A. The Kinase inactive (KI) mutant corresponds to LRRK1[D1409A]. 24 h post-transfection the cells were lysed and extracts (20 μg) from a duplicate experiment in which cells cultured in separate dishes were subjected to immunoblot analysis with the indicated antibodies (all at 1 µg/ml). Each lane represents cell extract obtained from a different replicate. The membranes were developed using the LI-COR Odyssey CLx Western Blot imaging system. Subsequent quantification presents the mean with error bars representing SEM. Similar results were obtained in 3 independent experiments. In SFig 5 data is shown that the double LRRK1[Y971F+K746G] mutant does not stimulate LRRK1 activity beyond the single mutants.

**Figure 6.**
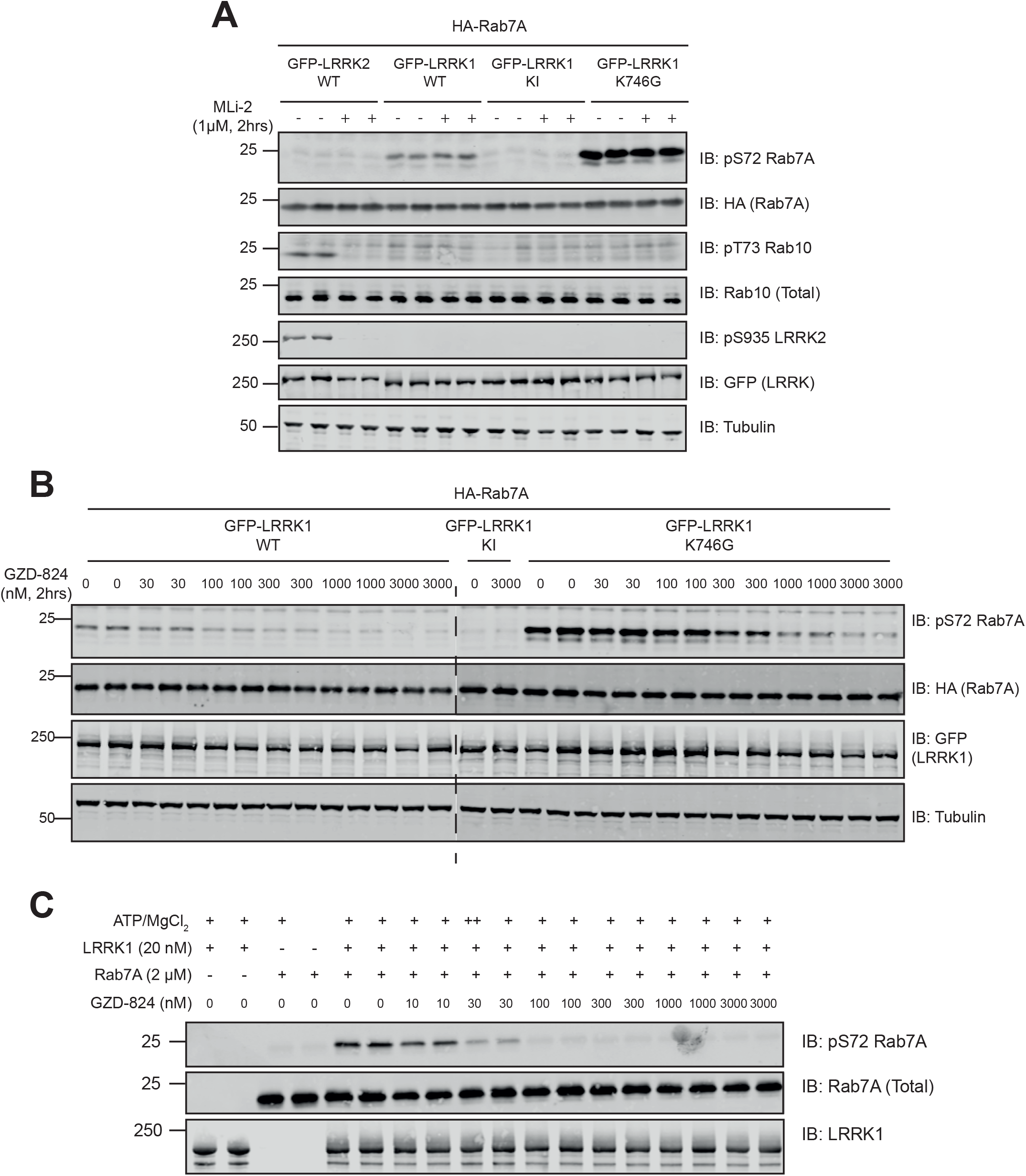
LRRK1 is inhibited by GZD-824 but not MLi-2. HEK293 cells were transiently transfected with the indicated plasmids encoding for wild type Rab7A and the indicated wild type and mutant forms of LRRK1 or wild type LRRK2. 24 h post-transfection, cells were treated with the indicated concentrations of MLi-2 (A) or GZD-824 (B) for 2 h. Cells were lysed and extracts (20 μg) from a duplicate experiment in which cells were cultured in separate dishes were subjected to immunoblot analysis with the indicated antibodies (all at 1 µg/ml). Each lane represents cell extract obtained from a different replicate. The membranes were developed using the LI-COR Odyssey CLx Western Blot imaging system. Similar results were obtained in 3 independent experiments. In SFig 6 we present the equivalent data undertaken with GSK2578215A, HG10-102-01, LRRK2-IN1 and iN04 inhibitors. (C) Wild type Rab7A was incubated in a phosphorylation reaction with recombinant LRRK1 as described in the materials and methods in the presence of the indicated concentrations GZD-824. Quantifications of experiments performed in (B) and (C) are presented in SFig 7. Reactions were terminated after 60 min with SDS-Sample buffer. Samples were subjected to immunoblot analysis with the indicated antibodies (all at 1 µg/ml). The membranes were developed using the LI-COR Odyssey CLx Western Blot imaging system.

We also analyzed the impact of two LRRK1 mutations (E929 truncation [9] and E1980A-FS [6]) reported to cause OSMD. The E1980A-FS mutation adds a 66 amino acid extension to the WD40 domain at the C-terminus (Fig 5B). These mutations block Rab7A phosphorylation in HEK293 cell overexpression assay (Fig 5D). We also tested 4 LRRK1 variants that have been reported in a SNP database (Supplementary Table 1) (https://www.ncbi.nlm.nih.gov/snp/; [53]) namely LRRK1[E929K], LRRK1[E929Q], LRRK1[Y971C] and LRRK1[I1412F]. Three of these mutations (LRRK1[E929K], LRRK1[E929Q] and LRRK1[I1412F]) induced a moderate, under 2-fold increase in pRab7A phosphorylation. The LRRK1[Y971C] variant, unlike the Y971F mutation, does not alter LRRK1-mediated Rab7A phosphorylation (Fig 5D).

### GZD-824 but not selective LRRK2 inhibitors inhibits LRRK1

As there was no robust assay for measuring LRRK1 activity, it was not known whether widely utilized LRRK2 inhibitors also inhibit LRRK1. We therefore tested whether some of the most widely used LRRK2 inhibitors impact LRRK1-mediated Rab7A phosphorylation. These studies revealed that none of the following inhibitors influenced Rab7A phosphorylation when tested in a HEK293 cell overexpression experiment when used at a 10-fold higher concentration than required to suppress LRRK2 activity: MLi-2 [44] (Fig 6A), GSK2578215A [54] (SFig 7A), HG10-102-01 [55] (SFig 7B), and LRRK2-IN1 [56] (SFig 7C),. All inhibitors ablated LRRK2-mediated phosphorylation of Rab10 at Thr73 and LRRK2 at Ser935 in parallel experiments. An earlier study reported that a compound identified form a virtual homology screen termed iN04 inhibited LRRK1, but no experimental data was presented to demonstrate this [57]. We found that iN04, when used at 5 μM had no impact on LRRK1-mediated Rab7A phosphorylation (SFig 7D).

The only kinase inhibitor that we could identify that suppressed LRRK1 activity was a compound originally identified as a poorly selective Abl tyrosine kinase inhibitor, termed GZD-824 [58]. This inhibitor has also been reported to target LRRK2 [59]. GZD-824 is a Type-2 kinase inhibitor that binds to the protein kinase domain, trapping them in the open, inactive conformation [59]. We found that GZD-824 potently inhibited wild type LRRK1-mediated Rab7A phosphorylation in HEK293 cells with an IC50 of ∼75 nM, and LRRK1[K746G] mutant with an IC50 of ∼275 nM (Fig 6B, SFig 8). GZD-824 also inhibited LRRK1 mediated phosphorylation of Rab7A an in vitro assay undertaken with 1 mM ATP with an IC50 value of ∼20 nM (Fig 6C, SFig 8).

### LRRK1 mediated Rab7A phosphorylation is not regulated by VPS35 or Rab29

The Parkinson’s causing VPS35[D620N] mutation markedly enhances LRRK2-mediated Rab protein phosphorylation [23]. To verify whether this mutation would also impact LRRK1, we studied the phosphorylation of Rab7A in previously described wild type and VPS35[D620N] knock-in MEFs [23]. This revealed that basal as well as PMA stimulated Rab7A phosphorylation were unaffected by the VPS35[D620N] mutation (Fig 7A). As expected, LRRK2 regulated Rab10 Thr73 phosphorylation was markedly increased by VPS35[D620N] (Fig 7A). Previous work has also demonstrated that LRRK2 interacts with Rab29 that is localized on the Golgi apparatus [21, 22]. LRRK2 is normally largely cytosolic, but when co-expressed with Rab29 becomes significantly relocalized to the Golgi (Fig 7B). LRRK1 is also largely cytosolic but does not relocalize to the Golgi when co-expressed with Rab29 in parallel studies (Fig 7B). Consistent with this data, recent work indicates that Rab29 interacts with the Armadillo domain in LRRK2 [60], which is absent in LRRK1 (Fig 5B).

**Figure 7.**
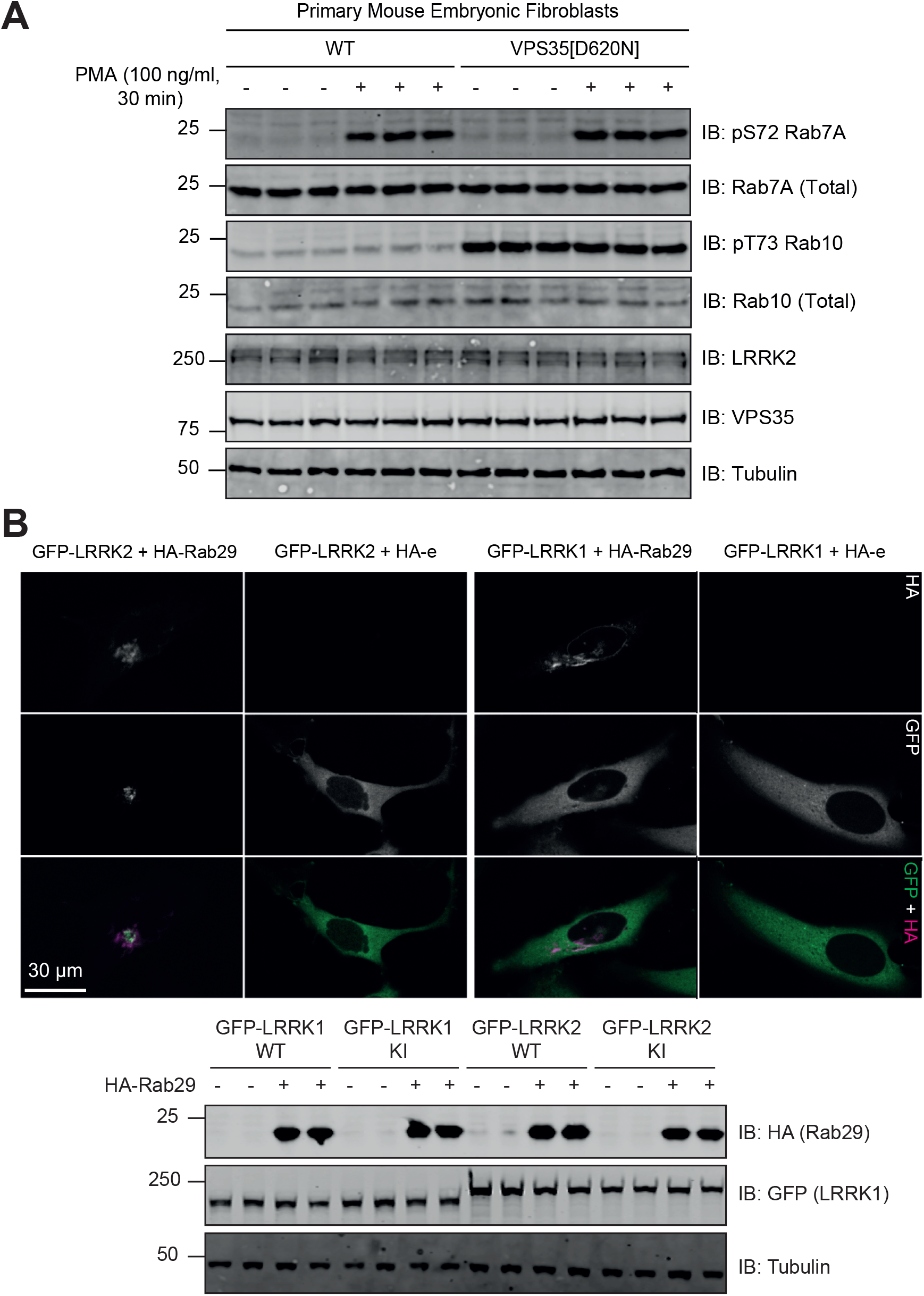
LRRK1 is not regulated by VPS35[D620N] mutation and overexpression of Rab29. (A) Wild type, and homozygous VPS35[D620N[knock-in primary MEFs were deprived of serum for 16 h and then incubated in the presence or absence of 100 ng/ml PMA for 30 min. Extracts (20 μg) from triplicate independent clones were subjected to immunoblot analysis with the indicated antibodies (all at 1 µg/ml). Each lane represents cell extract obtained from a different cell clone. The membranes were developed using the LI-COR Odyssey CLx Western Blot imaging system. Similar results were obtained in 3 independent experiments. (B, upper) HeLa cells were transfected with wild type GFP-LRRK1 or GFP-LRRK2 in the presence of absence of HA-Rab29. 24h post-transfection cells were fixed in 4% (by vol) paraformaldehyde and stained with mouse anti-HA. Scale bar represents 30 μm. (B, lower) As in (B, upper) except that cell are lysed and analyzed by immunoblot analysis as described in (A).

LRRK1 phosphorylation of Rab7A does not affect interaction with RILP Phosphorylation of Rab7A at Ser72 has been reported to enhance interaction with its downstream effector Rab-interacting lysosomal protein (RILP), in a manner dependent on Rab7A being bound to GTP, in its ‘active’ state [28]. In an attempt to confirm this, we co-expressed RILP with either wild type Rab7A or Rab7A[Q67L] (GTP trapped conformation) in the presence of LRRK1[K746G] to induce Rab7A phosphorylation. As a control we used the kinase inactive LRRK1[D1409A] mutant instead of the active LRRK1. Phos-tag analysis revealed that Rab7A and Rab7A[Q67L] were phosphorylated to a stoichiometry of ∼50% and ∼30% respectively (Fig 8, lower panel). Immunoprecipitation of RILP (Fig 8, upper panel) resulted in robust co-immunoprecipitation of Rab7A and vice versa immunoprecipitation of Rab7A (Fig 8, middle panel) resulted in co-immunoprecipitation of RILP. The co-immunoprecipitation of RILP with Rab7A was unaffected by Rab7A phosphorylation at Ser72, though there was a marginal decrease in co-immunoprecipitation with Rab7A[Q67L] in comparison to wild type (Fig 8).

**Figure 8:**
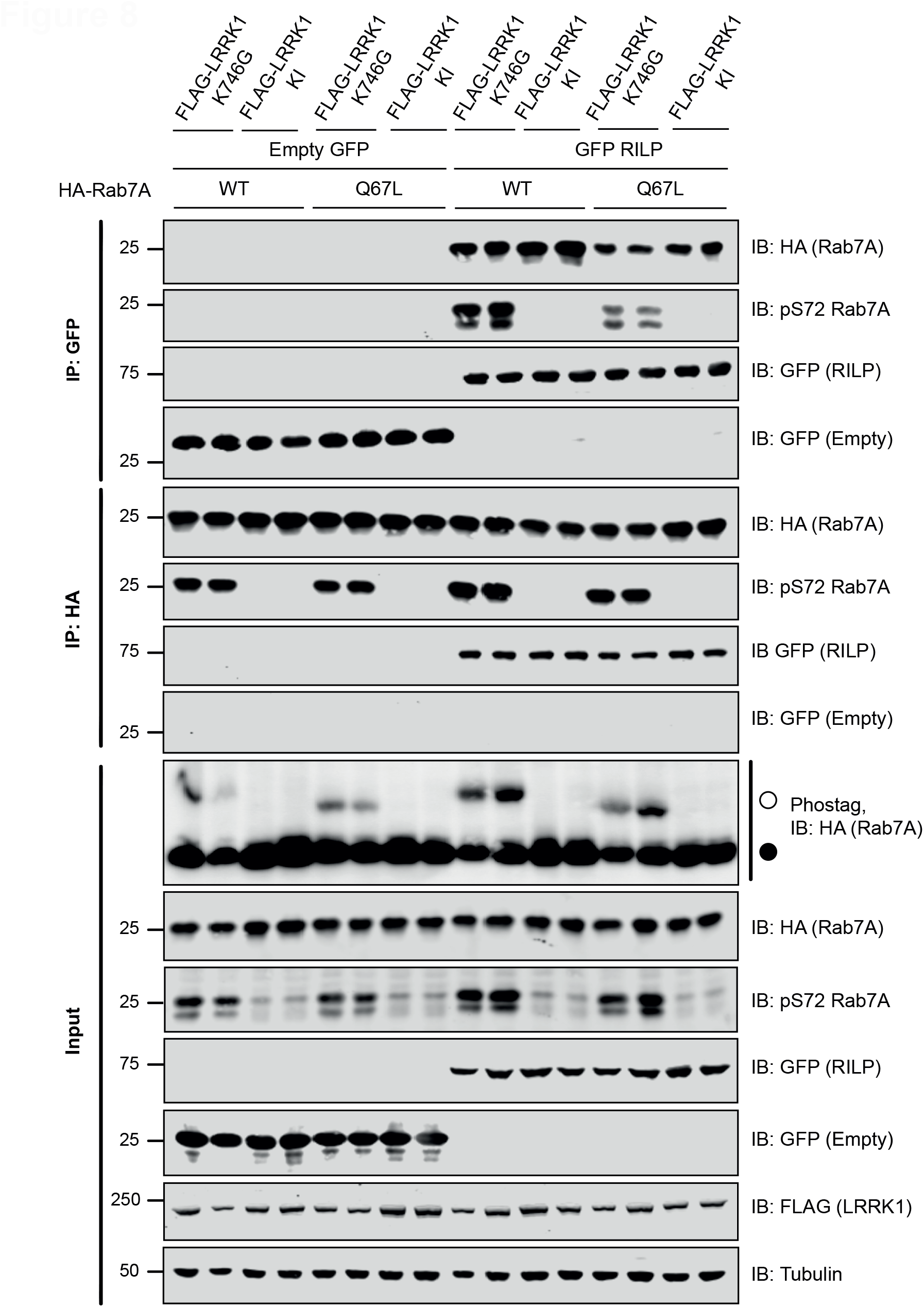
Phosphorylation of Rab7A does not affect interaction with RILP. HEK293 cells were transiently transfected with the indicated plasmids. Note that the Kinase inactive (KI) mutant corresponds to LRRK1[D1409A]. 24 h post-transfection cells were lysed either GFP-RILP (upper panel) or HA-Rab7A (middle panel) immunoprecipitated with GFP or HA antibody respectively. The immunoprecipitates as well as cell extracts (20 μg, lower panel) were subjected to immunoblot analysis with the indicated antibodies (all at 1 µg/ml). Each lane represents cell extract obtained from a different replicate. The membranes were developed using the LI-COR Odyssey CLx Western Blot imaging system.

### PPM1H, but not PPM1J or PPM1M, dephosphorylates Rab7A at Ser72

We next evaluated whether overexpression of PPM1H, a protein phosphatase that dephosphorylates LRRK2 phosphorylated Rab proteins [42], would also act on LRRK1 phosphorylated Rab7A. To test this, we transiently co-expressed Rab7A with LRRK1[K746G] to induce Rab7A phosphorylation and found that co-expression with wild type PPM1H, but not a catalytically inactive PPM1H[D153D] mutant [42], blocked Rab7A phosphorylation (Fig 9, upper panel). Overexpression of two other phosphatases most closely related to PPM1H namely PPM1J (Fig 9, middle panel) and PPM1M (Fig 9, lower panel) did not impact on LRRK1 mediated Rab7A phosphorylation.

**Figure 9.**
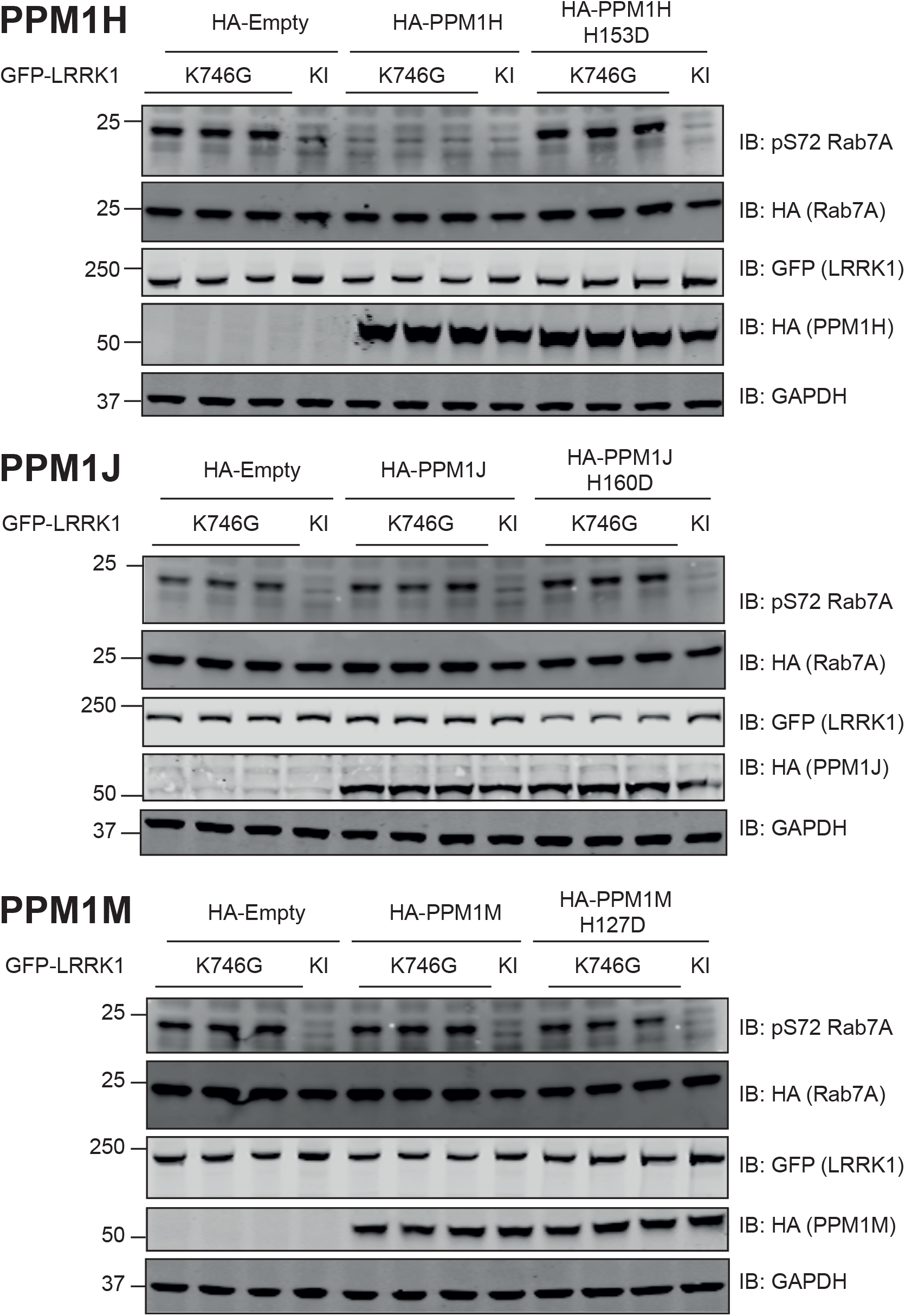
PP1MH, but not PPM1J or PPM1M, dephosphorylates Rab7A at Ser72. HEK293 cells were transiently transfected with the indicated plasmids encoding for wild type or catalytically inactive PPM1H (upper panel), PPM1J (middle panel) or PPM1M (lower panel), alongside LRRK1[K746G] or kinase inactive (KI) and wild type Rab7A. The Kinase inactive (KI) mutant corresponds to LRRK1[D1409A]. 24 h post-transfection the cells were lysed and extracts (20 μg) from a duplicate experiment in which cells cultured in separate dishes were subjected to immunoblot analysis with the indicated antibodies (all at 1 µg/ml). Each lane represents cell extract obtained from a different replicate. The membranes were developed using the LI-COR Odyssey CLx Western Blot imaging system.

## Discussion

Our data reveal that like LRRK2, LRRK1 functions as a Rab protein kinase. LRRK1 efficiently phosphorylates Rab7A at Ser72 within the Switch-II motif. This finding is also consistent with a recent study [28]. Ser72 lies in the equivalent position to the LRRK2 phosphorylation site in other Rab proteins (e.g Thr73 in Rab10). Our data indicate that LRRK1 is highly specific for Rab7A and does not phosphorylate Rab8A or Rab10 in vitro or in vivo. Conversely, LRRK2 does not phosphorylate Rab7A, consistent with previous work [15]. In future work it will be important to understand the molecular mechanism underlying the substrate specificity of LRRK1 and LRRK2, which could be addressed in on-going structural work [59, 61]. In vivo, co-localization of LRRK1 and LRRK2 with substrate Rab proteins on target membranes is also likely to be a key determinant in regulating Rab protein phosphorylation in vivo. How Rab proteins are brought together with LRRK1 and LRRK2 on cellular membranes is largely unknown. Recent work has revealed that localization of LRRK2 at a membrane is sufficient to activate kinase activity [62]. It would be interesting to establish whether this is also the case for LRRK1. LRRK2 phosphorylates a subset of Rab proteins (Rab1, Rab3, Rab8A, Rab8B, Rab10, Rab12, Rab29, Rab35 and Rab43) [15]. Our work has focused on Rab7A, Rab8A and Rab10, but it is possible that LRRK1 could phosphorylate other Rab proteins that we have not investigated. 50 Rab proteins possess a Ser/Thr residue at the equivalent position to Ser72 of Rab7A and thus far, at least 37 Rab proteins have been shown to be phosphorylated at this site when overexpressed in HEK293 cells [15]. It would be interesting to evaluate whether LRRK1 phosphorylates other Rab proteins in different tissues and cells especially osteoclasts.

There is strong evidence that LRRK1 is localized at endosomes especially following EGF receptor signalling where Rab7A is found [24, 26]. As mentioned in the introduction, previous work indicated that overexpression of LRRK1 induced phosphorylation of Rab7A at Ser72 and that this modulates EGF receptor trafficking [28]. We found that EGF stimulation in MEFs did not induce Rab7A phosphorylation (Fig 4A), despite activating the ERK signalling pathway (SFig 2). We have also studied Rab7A phosphorylation in HaCaT cells where EGF, but not PMA, stimulated Rab7A phosphorylation in a manner that was suppressed by siRNA knock-down of LRRK1 (SFig 4B). Further work will be required to decipher why EGF stimulates LRRK1 mediated Rab7A phosphorylation in HaCaT cells but not in MEFs and vice versa, why PMA only promotes Rab7A phosphorylation in MEFs. It would also be vital to study LRRK1 activity in osteoclasts given loss of LRRK1 activity in both humans and mice results in severe osteopetrosis [6, 7, 13]. How PMA induces phosphorylation of Rab7A in MEFs needs also to be further addressed. It is possible that the phorbol ester triggers the recruitment of LRRK1 to late endosomes where Rab7A is localized and that co-localization at this membrane compartment is sufficient to activate LRRK1 and drive Rab7A phosphorylation. As phosphorylation of Rab7A induced by phorbol ester is blocked with the general PKC inhibitor Go-6983, it is likely that PKC isoforms orchestrate LRRK1-mediated Rab7A phosphorylation in MEFs. Thus, it will be important in future work to study how PKC is linked to LRRK1, and whether this controls its activity, phosphorylation and/or cellular localization.

We have also developed robust in vitro and in vivo assays to interrogate LRRK1 kinase activity. We show that 4 widely used LRRK2 inhibitors do not inhibit LRRK1. Given the links of LRRK1 to osteopetrosis, inhibiting this enzyme would be undesirable. Furthermore, previous work has revealed that deleting both LRRK1 and LRRK2 in mice results in earlier mortality and age-dependent, selective neurodegeneration that is not observed in either of the single knock-out mice [63]. It would be important to verify that the different classes of LRRK2 inhibitors undergoing clinical trials, whose structures have not been revealed to date, are confirmed not to inhibit LRRK1. Our data suggest that the compound IN04, proposed through a virtual modelling screen with no experimental validation to be an LRRK1 inhibitor [34], does not significantly inhibit LRRK1 mediated Rab7A phosphorylation, at least at 5 μM, the concentration we tested.

Recent structural work indicates that the kinase domain of wild type LRRK2 is folded in an open, inactive state sometimes referred to as a Type-2 conformation [59]. This likely accounts for why the activity of wild type LRRK2 is relatively low compared to other active kinases. Mutations that hyperactivate LRRK2 are located within the ROC (R1441C/G), COR (Y1699C) and kinase domains (G2019S, I2020T). Mutations within the ROC and COR domains are thought to inhibit GTPase activity and promote the GTP bound conformation [64] which is thought to induce the kinase domain to adopt a closed active state-referred to as the Type 1 conformation [59]. The Parkinson’s mutations located within the LRRK2 kinase domain are reported to destabilize the Type-2 conformation in a manner that promotes LRRK2 to adopt its more closed Type-1 conformation [59, 65]. Our finding that LRRK1 is inhibited by the broadly selective GZD-824 inhibitor, a compound that is known to bind kinases in a Type 2 conformation [58], is consistent with wild type LRRK1 also adopting an open Type-2 conformation. Our observation that the LRRK1[K746G] mutant is ∼4-fold less potently inhibited by the Type-2 GZD-824 inhibitor than wild type LRRK1 (Fig 6C) is consistent with the LRRK1[K746G] mutant adopting a more active Type-1 conformation compared to wild type LRRK1, that would be expected to decrease affinity for a Type-2 kinase inhibitor.

Our finding that two mutations LRRK1[K746G] (equivalent to LRRK2[R1441G]) and LRRK1[I1412T] (equivalent to LRRK2[I2020T]) promoted Rab7A phosphorylation also indicates that LRRK1 could be regulated similarly to LRRK2. The LRRK2[R1441G] mutation in the ROC domain increases LRRK2 mediated phosphorylation of Rab10, 3 to 4-fold. The LRRK2[R1441G] mutation also enhances affinity of LRRK2 towards its upstream activator, Rab29 [21]. The LRRK1[K746G] mutation also induces robust ∼5-fold increase in Rab7A phosphorylation. It would be interesting to investigate whether the K746G mutation also suppresses GTPase activity, enhancing GTP binding. The LRRK2[I2020T] mutation increases Rab10 phosphorylation ∼2-fold, to a similar extent as the LRRK1[I1412T] mutation promotes Rab7A phosphorylation. There are some differences between LRRK1 and LRRK2, as the LRRK1[F1022C] (equivalent to LRRK2[Y1699C]) and LRRK1[G1411S] (equivalent to LRRK2[G2019S]) do not enhance LRRK1-mediated phosphorylation of Rab7A. We observed that the previously reported LRRK1[Y971F] activating mutation [27, 28] within the COR dimerization domain, promoted Rab7A phosphorylation to the same extent as the LRRK1[K746G] mutation. Previous work suggested that this mutation exerts its effect by blocking an inhibitory phosphorylation by the EGF receptor [27, 28]. We have not stimulated cells with EGF, so it is unlikely that this site would be phosphorylated by the EGF receptor to a high stoichiometry. Moreover, we find that the LRRK1[Y971C] variant [53] that would also block EGF receptor phosphorylation is not activated (Fig 5D). It would be worth investigating whether the LRRK1[Y971F] mutation promoted LRRK1 activation by inhibiting GTPase activity thereby enhancing the GTP bound conformation of LRRK1. This might also account for why combining the K746G and Y971F mutations does not further enhance LRRK1 activity (SFig 6), if both of these mutations are working the same way to promote a GTP bound conformation. In future work it would be interesting to generate a knock-in cells/mouse to explore the physiological impact that LRRK1 activating mutations have.

Mutations that delete the C-terminal 8 residues of LRRK2 ablate kinase activity [66]. Recent structural work reveals that the C-terminal 28 residues of LRRK2 form an elongated *α*-helix that extends along the entire kinase domain, interacting with both its C-and N-lobes that likely stabilize the kinase domain and may form binding platforms for other LRRK2 regulators [59]. Several autosomal recessive variants that induce frameshift or truncating mutations impact the equivalent C-terminal residues of LRRK1 and cause OSMD [6-9]. We have found that one of these mutations, namely E1980A-FS, impacts the C-terminal 66 residues and inactivates LRRK1 (Fig 5D). This would be consistent with the C-terminal motif of LRRK1 playing a similar key role in controlling kinase activity. It is likely that all LRRK1 mutations that have been linked to OSMD inhibit LRRK1 kinase activity.

Previous work indicated that Rab7A phosphorylation is mediated by the PINK1/TBK1 signalling pathways in experiments undertaken in HeLa cells overexpressing Parkin [48]. In wild type MEFs, we found that activation of the PINK1 or TBK1 kinases failed to induce significant phosphorylation of Rab7A, relative to PMA stimulation. Previous work studying Rab7A phosphorylation by PINK1/TBK1 pathways was undertaken using mass spectrometry [48] that is more sensitive than immunoblotting approaches used in our study. We cannot rule out that the PINK1/TBK1 system induces low stoichiometry phosphorylation of Rab7A that we are unable to detect by immunoblotting. Previous work suggested that the PTEN phosphatase is capable of dephosphorylating Rab7A that is phosphorylated at Ser72 as well as Tyr183, which were reported necessary for GDP dissociation inhibitor dependent recruitment of Rab7 onto late endosomes and subsequent maturation [67]. We have not tested whether PTEN can dephosphorylate Rab7A.

Our data show that the PPM1H protein phosphatase previously shown to dephosphorylate Rab proteins phosphorylated by LRRK2 [42], was also capable of dephosphorylating Rab7A phosphorylated at Ser72 (Fig 9, upper panel). This indicates that PPM1H may not be specific towards dephosphorylating LRRK2 phosphorylated Rab proteins but may display broader specificity towards a wider group of switch-II motif phosphorylated Rab proteins. The related PPM1J and PPM1H phosphatases did not dephosphorylate Rab7A in parallel experiments (Fig 9, middle and lower panels).

In future work it would be also be important to probe the functional consequences of Rab7A Ser72 phosphorylation. In a previous study where stoichiometry of Rab7A was not verified, it was reported that overexpression of LRRK1[Y971F] mutant enhanced binding of Rab7A to one of its well characterized receptors termed RILP [28]. In our experiments we found that LRRK1 phosphorylation of Rab7A to a stoichiometry of ∼50%, did not noticeably impact levels of RILP that co-immunoprecipitated with Rab7A (Fig 8). Inspection of the crystal structure of Rab7A in complexes with RILP [68], reveals that RILP binds quite differently to Rab7A compared to RILPL2 binding to Thr72 phosphorylated Rab8A [19]. The conserved Arg132 residue in RILPL2 that binds to phosphorylated Thr72 in Rab8A is higher up in RILP and not part of interface of the complex with Rab7A. Ser72 in Rab7A interacts with Glu249 in RILP, but there is no obvious nearby basic residue that would explain how phosphorylation of Ser72 could enhance binding to RILP. Our analysis indicates that the phosphorylated Ser72 could be accommodated in the RILP complex. In future work it would be important to undertake more quantitative analysis of the interaction of RILP with phosphorylated versus dephosphorylated Rab7A. By analogy to the LRRK2 pathway, phosphorylation of Rab8A and Rab10 triggers interaction with a set of effectors (RILPL1, RILPL2, JIP3 and JIP4) that possess an RH2 motif that acts as a phospho-recognition domain for the LRRK2 phosphorylated Rab proteins [15, 19]. It is possible that much of the downstream physiology controlled by LRRK2 such as ciliogenesis [20, 69-71], is mediated through these phosphorylation-specific effector binding to phosphorylated Rab proteins. It would therefore be interesting to explore whether an equivalent set of phosphorylation-specific effectors exists for LRRK1 phosphorylated Rab7A and to dissect their physiological roles.

## Acknowledgements

We thank Odetta Antico for providing PINK1 reagents and advise how to study this pathway. Kuldip Dave, The Michael J. Fox Foundation for generating and depositing the LRRK1 KO mice at the JAX laboratories, the excellent technical support of the MRC-Protein Phosphorylation and Ubiquitylation Unit (PPU) DNA Sequencing Service (coordinated by Gary Hunter), the MRC-PPU tissue culture team (coordinated by Edwin Allen), MRC PPU Reagents and Services antibody and protein purification teams (coordinated by Hilary McLauchlan and James Hastie). We also thank Dr. Renata F. Soares from MRC-PPU mass spectrometry facility with her help in the maintenance of the LC and mass spectrometer for the analysis.

## Funding

This work was supported by the Michael J. Fox Foundation for Parkinson’s research [grant number 17298 (D.R.A.)] and [grant number 6986 (D.R.A.)], the Medical Research Council [grant number MC_UU_12016/2 (D.R.A.)] and the pharmaceutical companies supporting the Division of Signal Transduction Therapy Unit (Boehringer-Ingelheim, GlaxoSmithKline, Merck KGaA -to D.R.A.).

## Authors Contribution

A. U. M. designed, executed experiments in Figures 2B, 3A, 3D, 5A-D, 6A-C, 9, S6, S7A-D, S8 and wrote the manuscript with D. R. A. A. K. designed and executed experiments in Figures 4A, 4D-F, S2-S5. F. T. was responsible for the generation of LRRK1 WT and KO MEFs, in addition to designing and executing experiments in figures 1B, 2A, 3B, 3C, 4B and 4C. Also performed immunoblotting of total Rab protein levels in figure 1A. R. S. N. performed PRM assay to look at levels of phospho-Rab proteins in LRRK1 WT and KO MEFs in Figure 1A. S. M. and S.K. planed and produced and purified recombinant LRRK1 protein and provided figure S1. R. G. performed Edman degradation sequencing with F. T. to identify phosphorylation sites on Rab7A and also contributed figure 3C. N.K.P. oversaw and coordinated the generation of the of MJF-38 rabbit polyclonal pS72 Rab7A antibody. P.L undertook all work associated with characterization of the potency and specificity of the MJF-38 rabbit polyclonal pS72 Rab7A antibody. M. T. designed and executed experiment in figure 7A, E. P. designed and executed experiment in figure 7B, and P. P. designed and executed experiment in figure 8. M. D. was involved in cloning, production and purification of recombinant Rab7A[WT], Rab7A[S72A] and Rab10[WT] proteins. S. W. and R. T. were responsible for the production of cDNA clones used for overexpression experiments. D. R. A. assisted with experimental design, formal analysis, supervision, funding acquisition, and writing the manuscript with A. U. M.

## Data and Reagents Availability

All cDNA clones, antibodies, proteins and other reagents generated at the University of Dundee can be requested by contacting the authors or via our website (https://mrcppureagents.dundee.ac.uk/). All other data can be requested by contacting the authors. The Mass Spectrometry data is being submitted to the PRIDE repository.

## Figure Legends

**Supplementary Figure S1:**
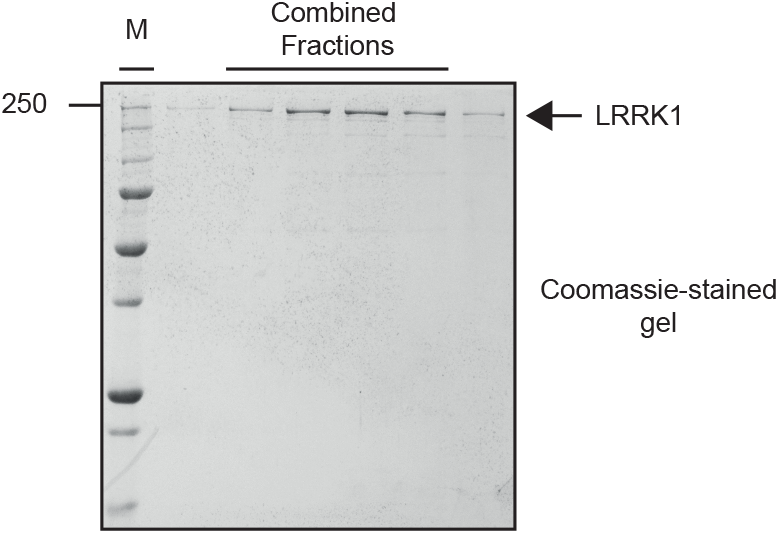
Coomassie gel of the purified recombinant LRRK1. 20 μl of the final gel filtration purification encompassing the peak of LRRK1 protein were analyzed by SDS-polyacrylamide electrophoresis and stained with Coomassie Blue. The fractions that were combined are indicated with a solid line.

**Supplementary Figure S2.**
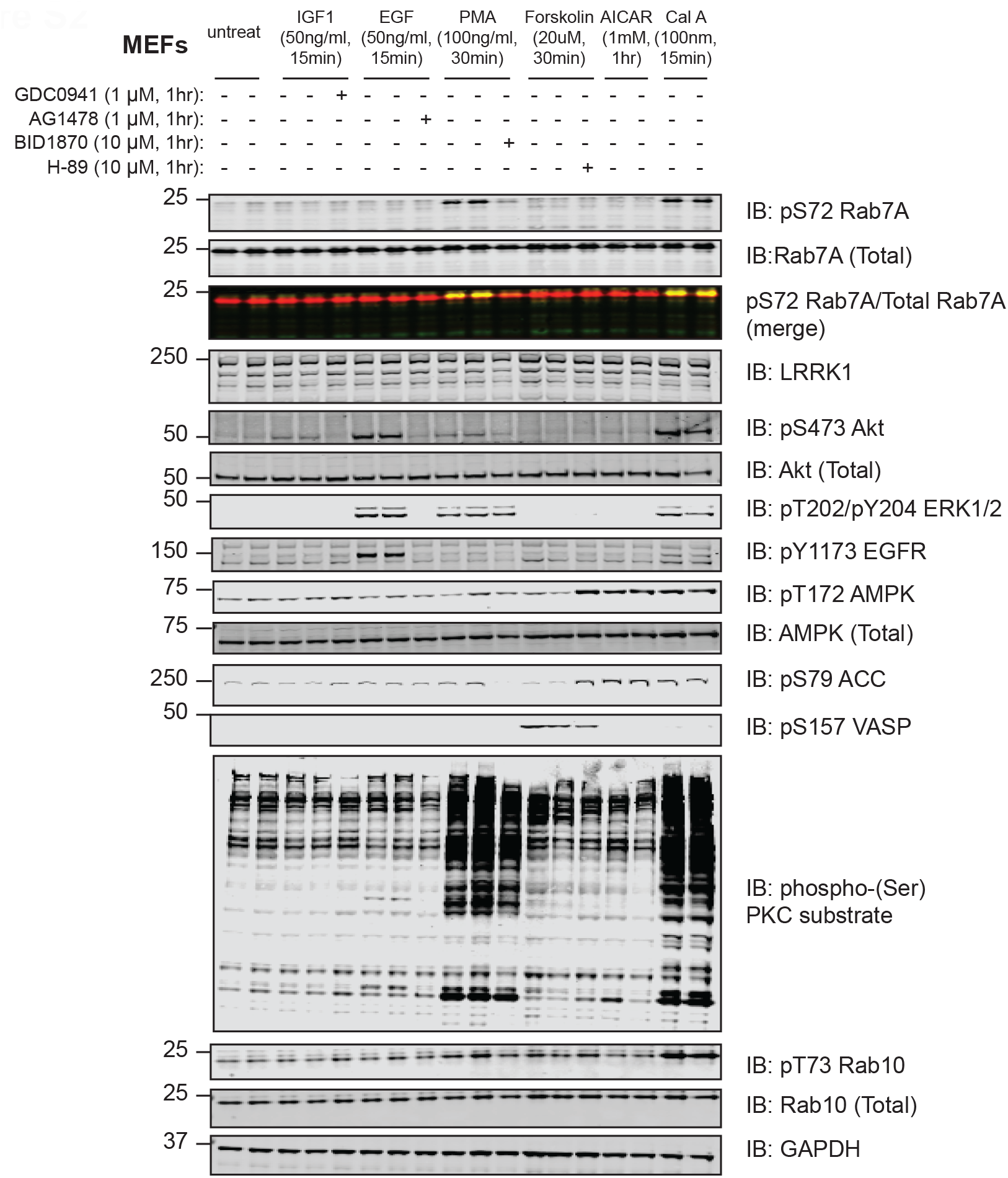
PMA and calyculin A stimulates Rab7A phosphorylation in wild type MEFs. Wild type MEFs were deprived of serum for 16 h and then incubated for 1 h in the presence or absence of the dose of indicated inhibitor. Cells were then stimulated with the dose of indicated agonists for the times indicated. Extracts (20 μg) from duplicate experiment were subjected to immunoblot analysis with the indicated antibodies (all at 1 µg/ml). Each lane represents cell extract obtained from a different cell dish. The membranes were developed using the LI-COR Odyssey CLx Western Blot imaging system.

**Supplementary Figure S3.**
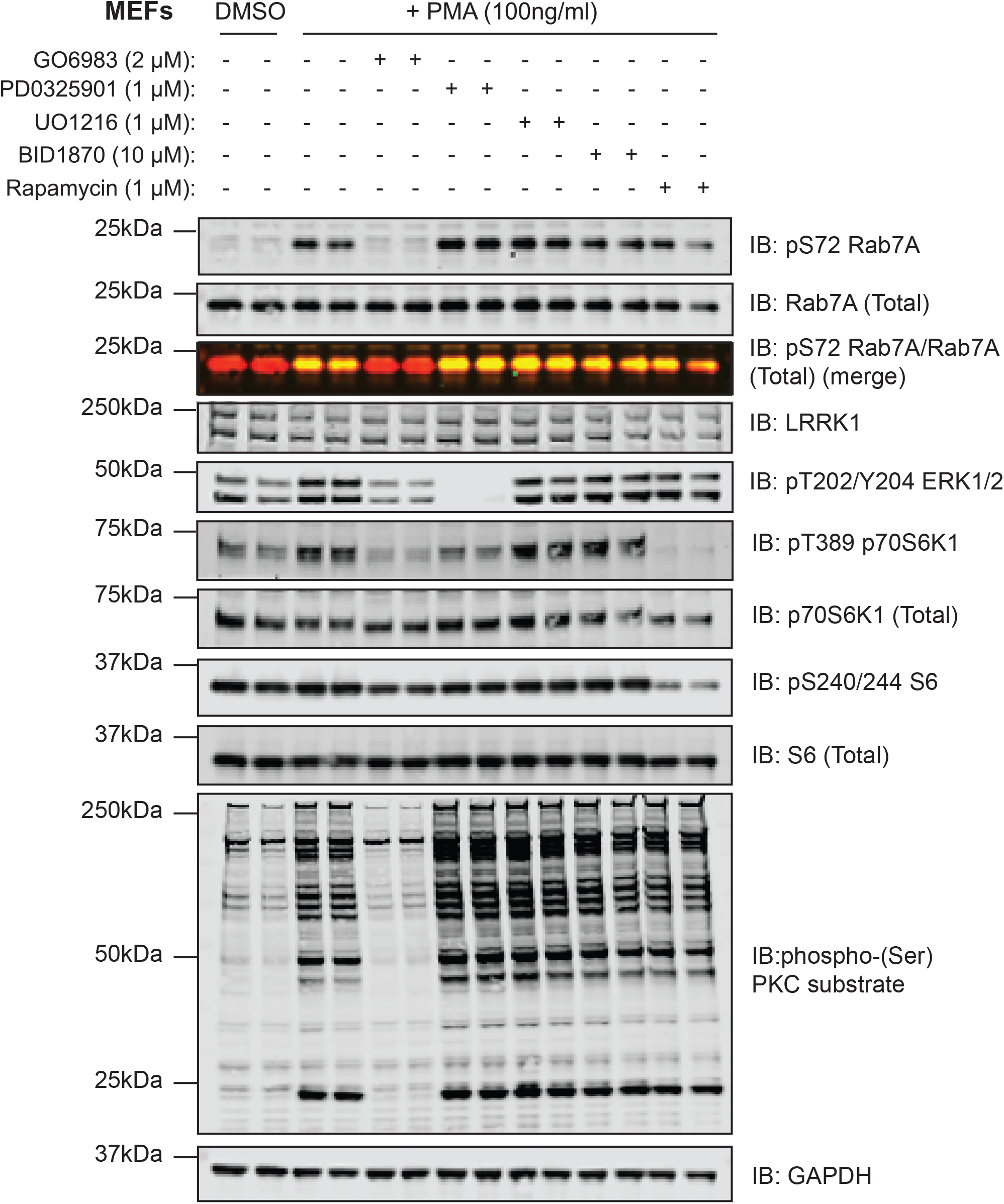
Rab7A phosphorylation induced by PMA is blocked by GO6983 PKC inhibittor. Wild type MEFs were deprived of serum for 16 h and then incubated for 1 h in the presence or absence of the dose of indicated inhibitor. Cells were then stimulated with PMA (100 ng/ml) for 30 min. Extracts (20 μg) from duplicate experiment were subjected to immunoblot analysis with the indicated antibodies (all at 1 µg/ml). Each lane represents cell extract obtained from a different cell dish. The membranes were developed using the LI-COR Odyssey CLx Western Blot imaging system.

**Supplementary Figure S4.**
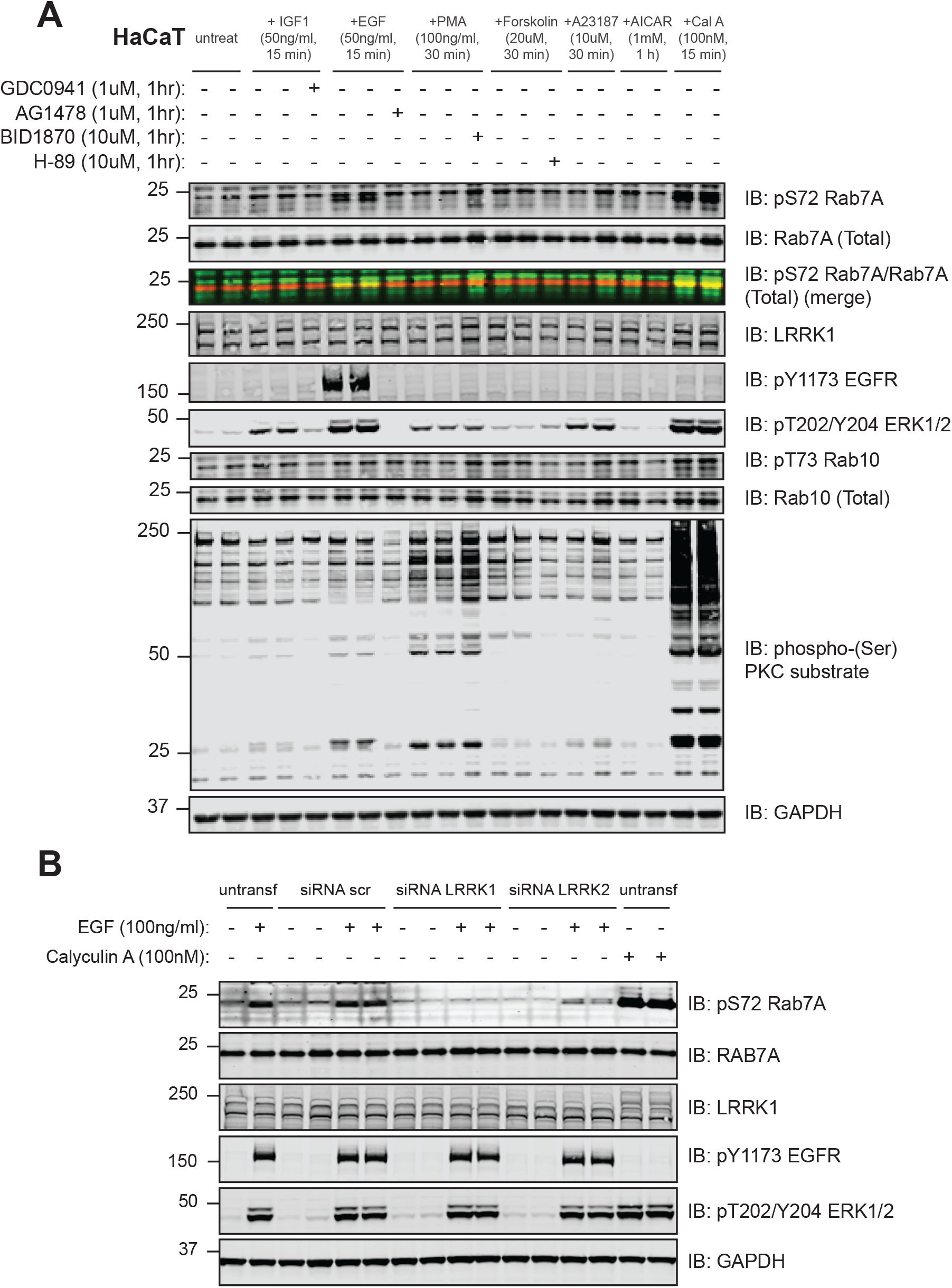
EGF-stimulation induces Rab7A phosphorylation at Ser72 by LRRK1 in HaCaT cells. (A) Wild type HaCat cells (human immortal keratinocytes) were deprived of serum for 16 h and then incubated for 1 h in the presence or absence of the dose of indicated inhibitor. Cells were then stimulated with the dose of indicated agonists for the times indicated. Extracts (20 μg) from duplicate experiment were subjected to immunoblot analysis with the indicated antibodies (all at 1 µg/ml). Each lane represents cell extract obtained from a different cell dish. The membranes were developed using the LI-COR Odyssey CLx Western Blot imaging system. (B) 2×10^5^ HaCaT cells/well were seeded in 6-well plates and after 24h transfected with 10nM of siRNAs against LRRK1, siRNA with scrambled sequence (scr) or left untransfected. After 60h, cells were serum starved overnight and then stimulated with EGF (50 ng/ml) for 15 min. Serum-starved, untransfected cells treated with Calyculin A (100 nM, 5 min) served as control. Immunoblots were developed using LI-COR Odyssey CLx Western Blot imaging system with the indicated antibodies at 1 μg/ml concentration.

**Supplementary Figure S5.**
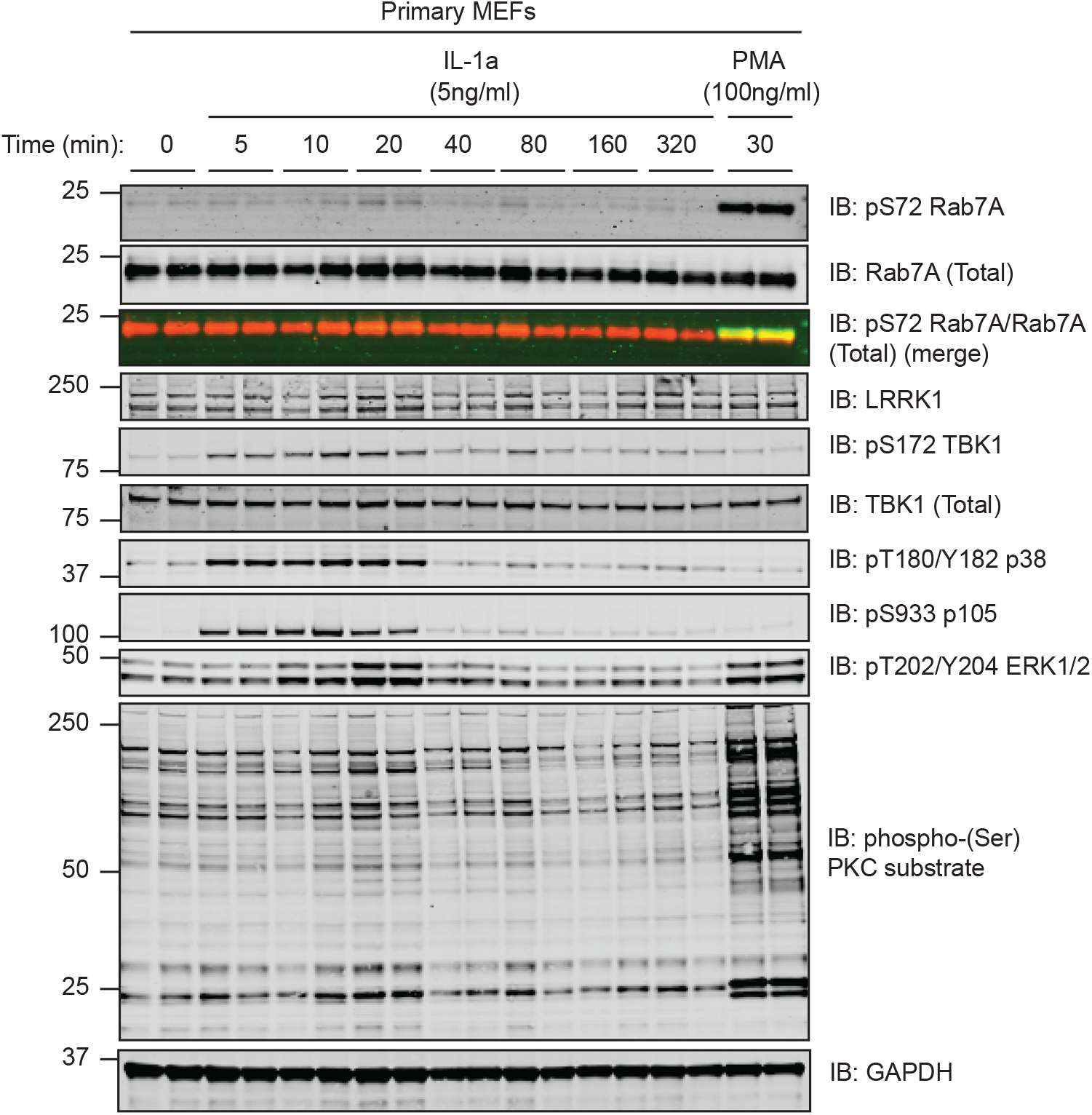
TBK1 activation peaks after 20 mins of IL-1A treatment in wild type MEFs. Wild type MEFs were deprived of serum for 16 h and then treated with IL-1A at the indicated time points. Extracts (20 μg) from duplicate experiments were subjected to immunoblot analysis with the indicated antibodies (all at 1 µg/ml). Each lane represents cell extract obtained from a different cell dish. The membranes were developed using the LI-COR Odyssey CLx Western Blot imaging system.

**Supplementary Figure S6:**
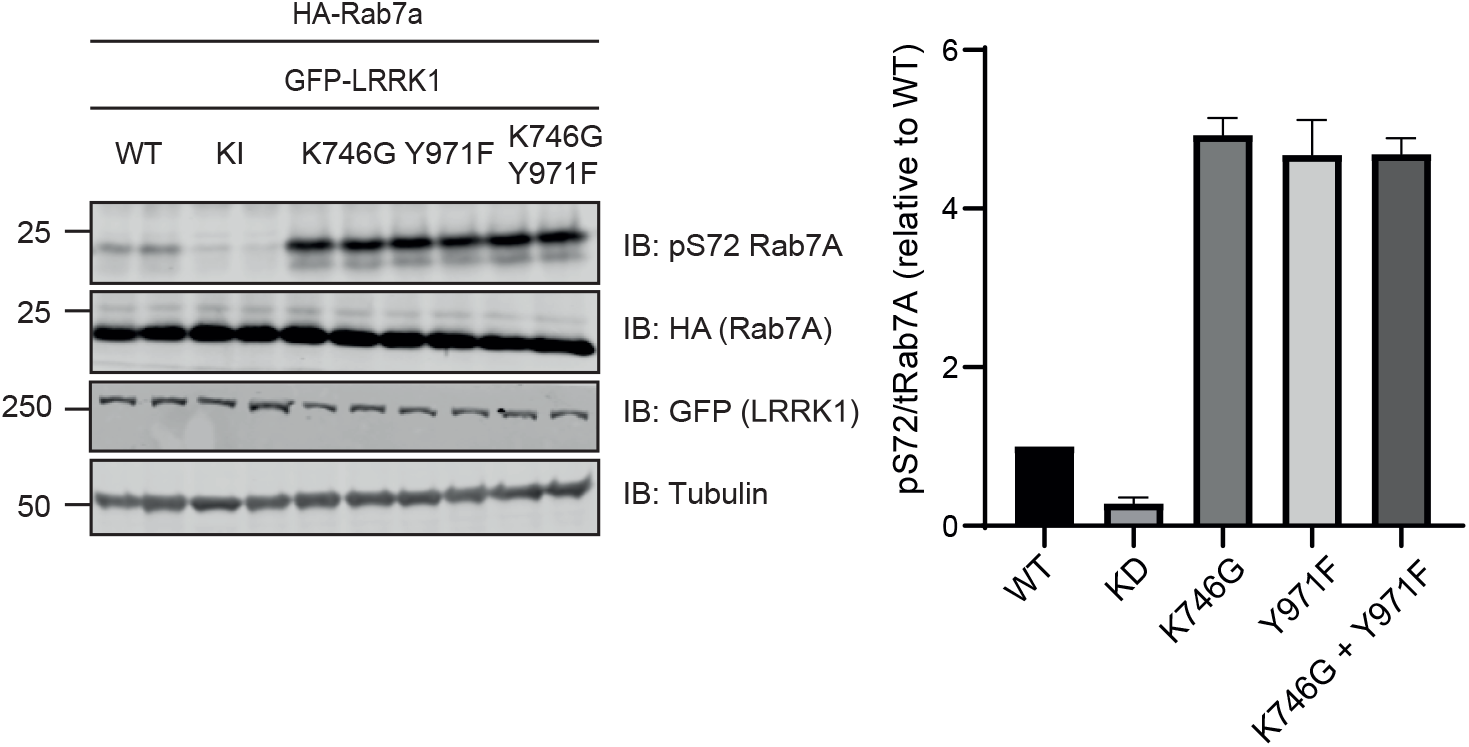
The LRRK1[K746G+Y971F] double mutant does not further activate LRRK1. HEK293 cells were transiently transfected with the indicated plasmids encoding for wild type and mutant of LRRK1 and wild type Rab7A. Note that the Kinase inactive (KI) mutant corresponds to LRRK1[D1409A]. 24 h post-transfections the cells were lysed and extracts (20 μg) from a duplicate experiment in which cells cultured in separate dishes were subjected to immunoblot analysis with the indicated antibodies (all at 1 µg/ml). Each lane represents cell extract obtained from a different replicate. The membranes were developed using the LI-COR Odyssey CLx Western Blot imaging system

**Supplementary Figure S7.**
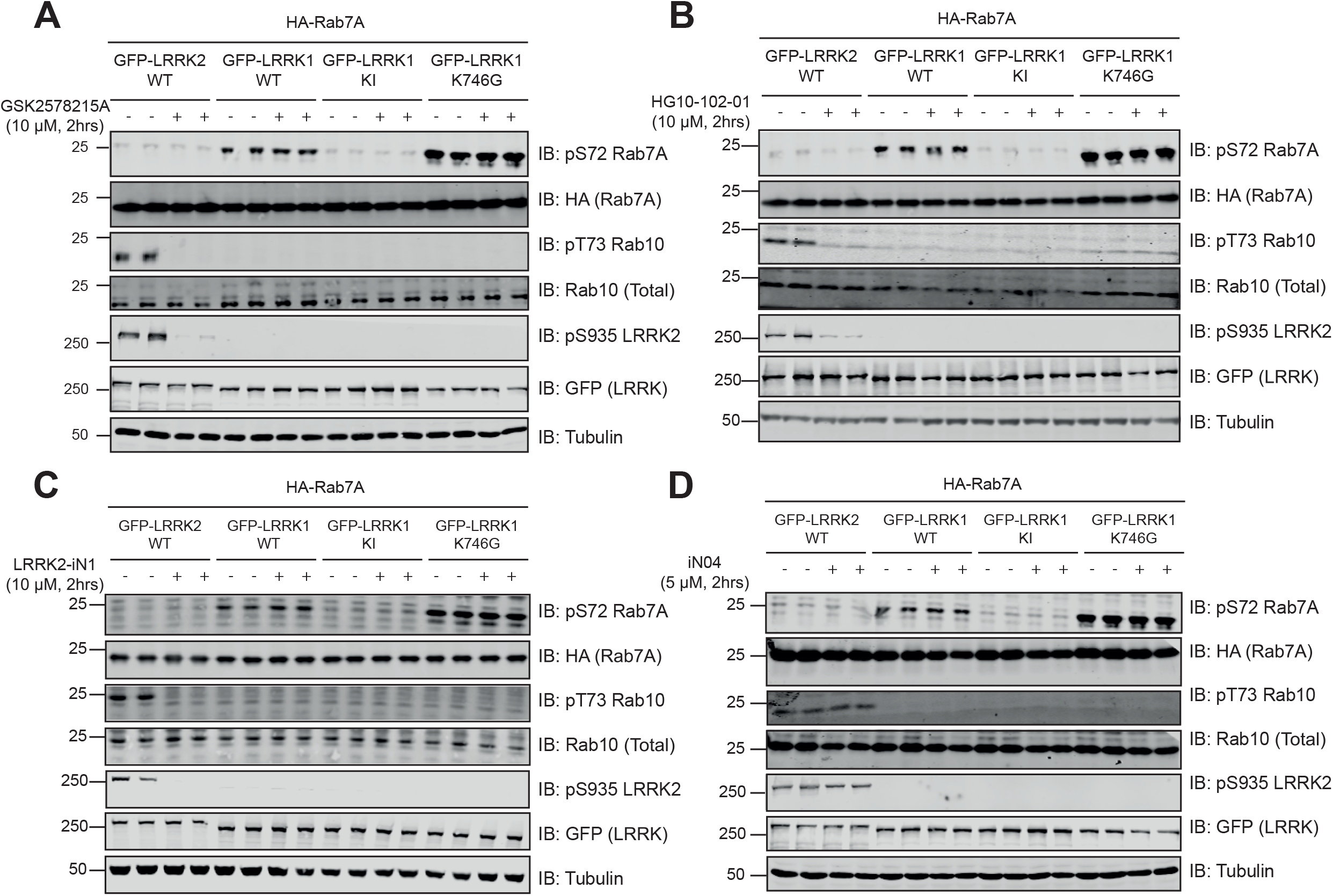
GSK2578215A, HG10-102-01, LRRK2-IN1 and iN04 do not inhibit LRRK1. HEK293 cells were transiently transfected with the indicated plasmids encoding for wild type Rab7A and the indicated wild type and mutant forms of LRRK1 or wild type LRRK2. 24 h post-transfections the cell were treated with the indicated concentrations of GSK2578215A (A), HG10-102-01 (B), LRRK2-IN1 (C) and iN04 (D) for 2 h. Cells were lysed and extracts (20 μg) from a duplicate experiment in which cells were cultured in separate dishes were subjected to immunoblot analysis with the indicated antibodies (all at 1 µg/ml). Each lane represents cell extract obtained from a different replicate. The membranes were developed using the LI-COR Odyssey CLx Western Blot imaging system. Similar results were obtained in 3 independent experiments.

**Supplementary Figure S8.**
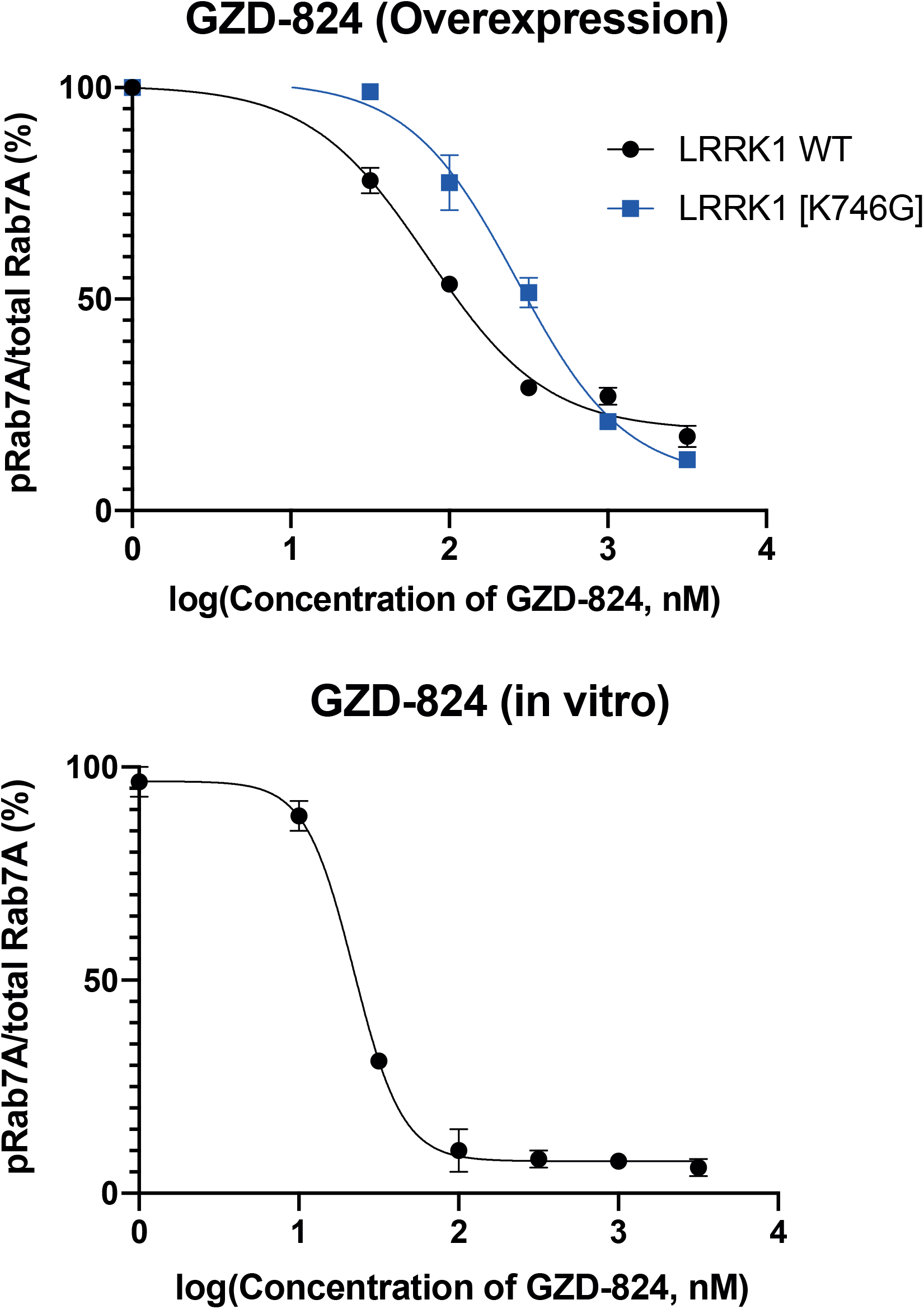
GZD-824 inhibits LRRK1 when overexpressed and *in vitro*. Quantification of GZD-824 dose-response experiments from Fig 6B (Upper) and 6C (Lower). Results are presented with log (Concentration of GZD-824, nM) on X-axis and pRab7A/total Rab7A relative to DMSO-treated control on Y-axis.

**Supplementary Table 1.**
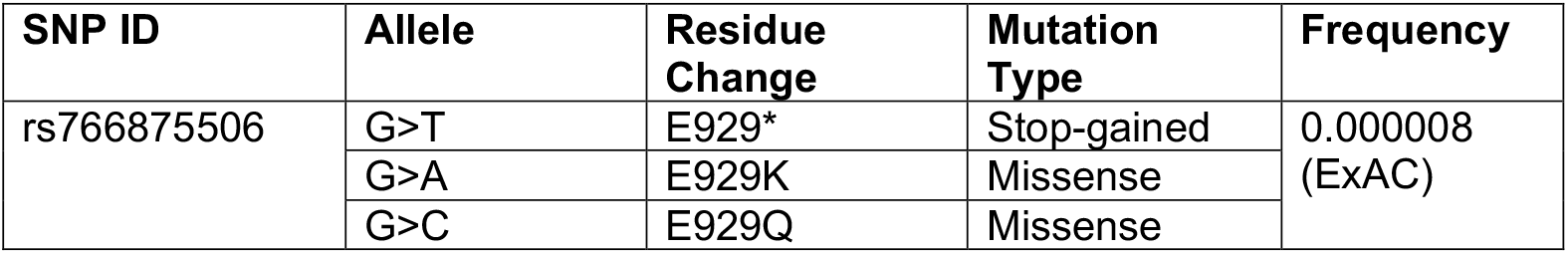
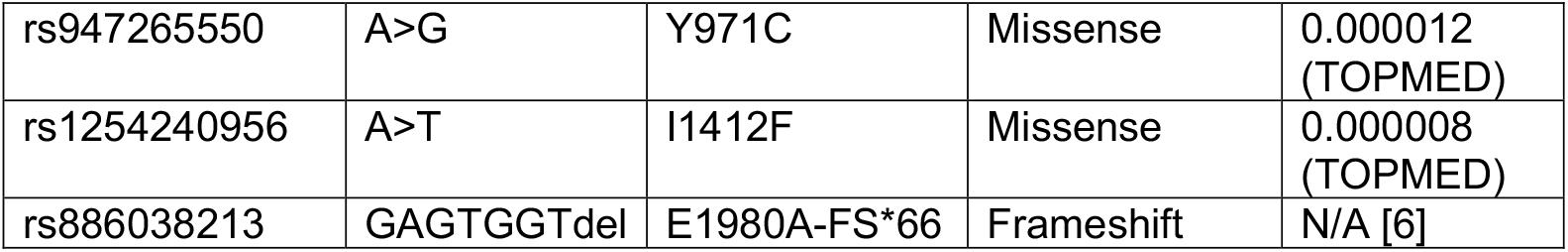
Single Nucleotide Polymorphisms in LRRK1.

**Supplementary File 1**.

The excel sheet contains the phospho Rab peptide sequences, the corresponding heavy and light m/z, their charge states and scheduled retention times used for targeted Mass spectrometry analysis of LRRK1 KO and LRRK1 wt MEFS.

## Notes

### Competing Interest Statement

The authors have declared no competing interest.

### Summary of Updates

Due to a technical error on submitting this manuscript one of the authors of the paper was inadvertently omitted on the bioRxiv author page (not on the PDF version of the manuscript). We have now corrected this issue and we apologise for not noticing this error when we made the original submission

